# The Mesodiencephalic Junction as a Central Hub for Cerebro-Cerebellar Communication

**DOI:** 10.1101/2021.02.23.432495

**Authors:** Xiaolu Wang, Manuele Novello, Zhenyu Gao, Tom J.H. Ruigrok, Chris I. De Zeeuw

## Abstract

Most studies investigating the impact of cerebral cortex (CC) onto the cerebellum highlight the role of the pontine mossy fibre system. However, cerebro-cerebellar communication may also be mediated by the olivary climbing fibres via a hub in the mesodiencephalic junction (MDJ). Here, we show that rostromedial and caudal parts of mouse CC predominantly project to the principal olive via the rostroventral MDJ and that more rostrolateral CC regions prominently project to the rostral medial accessory olive via the caudodorsal MDJ. Moreover, transneuronal tracing results show that the cerebellar nuclei innervate the olivary-projecting neurons in the MDJ that receive input from CC, and that they adhere to the same topographical relations. By unravelling these topographic and dense, mono- and disynaptic projections from the CC through the MDJ and inferior olive to the cerebellum, this work establishes that cerebro-cerebellar communication can be mediated by both the mossy fibre and climbing fibre system.

## Introduction

Cerebro-cerebellar communication can facilitate various forms of sensorimotor control (Popa et al., 2013, Hamada et al., 2012, Proville et al., 2014, Brown and Raman, 2018, Akkal et al., 2007, Quartarone et al., 2020) and cognitive processing (D’Mello and Stoodley, 2015, Depping et al., 2018, Olivito et al., 2017, Igelström et al., 2017, Gao et al., 2018, Crippa et al., 2016, Brissenden and Somers, 2019). For example, it is required for active exploration during online coordination (Onuki et al., 2015, Lindeman et al., 2021), as well as for motor planning, and action perception during social cognition (Abdelgabar et al., 2019, Gao et al., 2018, Chabrol et al., 2019). Cerebro-cerebellar communication is organized in multiple reciprocal loops. The information route from the cerebellum to the cerebral cortex (CC) includes either a direct connection from the cerebellar nuclei (CN) onto various subnuclei of the thalamus (Rispal-Padel and Grangetto, 1977, Shinoda et al., 1993, Amino et al., 2001), and/or a more indirect connection from the CN to prethalamic hubs like the zona incerta (Teune et al., 2000, Mitrofanis and deFonseka, 2001). Instead, information flow from the CC to the cerebellum is massively relayed via the pontine nuclei (Legg et al., 1989, Glickstein et al., 1985, Leergaard, 2003, Brodal, 1978). The ponto-cerebellar connection provides mossy fibres to the granule cells in the cerebellar cortex and collaterals to neurons in the CN (Henschke and Pakan, 2020, Biswas et al., 2019, Ruigrok, 2011), which is instrumental in planning and coordinating fine precision movements of the forelimbs (Wagner et al., 2019, Guo et al., 2020).

In addition to the mossy fibre route, CC may also control cerebellar processing via their climbing fibre input, which is derived from the inferior olive (IO) (Saint-Cyr, 1983, Burman et al., 2000, Ramnani, 2006, Kubo et al., 2018). Whereas decades of neuroanatomical research have elucidated the intricate topographical relationship of the cortico-ponto-cerebellar connections (Leergaard, 2003, Legg et al., 1989, Glickstein et al., 1985, Henschke and Pakan, 2020, Brodal, 1978), much less is known about those of the cortico-olivocerebellar system. Direct cerebro-olivary connections are sparse (Berkley and Worden, 1978, Saint-Cyr, 1983) and the indirect connections are diverse, presumably including various secondary intermediaries in both the lower and higher brainstem (Bull et al., 1990, Berkley et al., 1986). Of particular interest in this respect is the mesodiencephalic junction (MDJ), as this area provides a particularly dense projection to the IO (Mabuchi and Kusama, 1970, Linauts and Martin, 1978, Nakamura et al., 1983a, Swenson and Castro, 1983, Leichnetz et al., 1984, Onodera, 1984, Rutherford et al., 1984, de Zeeuw et al., 1989, Kubo et al., 2018). This area around the fasciculus retroflexus (FR) includes, amongst others, the medial accessory nucleus of Bechterew, the nucleus of Darkschewitsch, and it targets particularly the principal olive and the rostral medial accessory olive (Figure 1 - figure supplement 1) (Onodera, 1984, Saint-Cyr, 1987, de Zeeuw et al., 1989, Voogd and Ruigrok, 2004, De Zeeuw and Ruigrok, 1994). Next to an input from the CC, the MDJ also receives input from the CN (Saint-Cyr, 1987, Teune et al., 2000, Bentivoglio and Kuypers, 1982, Rutherford et al., 1989, Gonzalo-Ruiz et al., 1990, De Zeeuw and Ruigrok, 1994), raising the possibility that CC and CN conjunctively control olivary activity via this central hub in the brainstem. Despite all the evidence for its central strategic position in cerebro-cerebellar communication, several critical questions on the input-output relations of the MDJ remain to be elucidated. It is still unclear which olivary subnuclei receive inputs from which parts of the CC via which parts of the MDJ. In addition, it is also unclear whether the MDJ neurons that do receive input from the CC and project to the IO also receive a direct input from the CN, and if they do, from which parts of the CN. Moreover, in addition to these specific questions on the topography of the connections of MDJ neurons, it is also unclear how dense they are.

Here, we addressed these questions in mice by exploiting classical and advanced, retrograde and anterograde tracing approaches as well as by analyzing mesoscale cases of cortical anterograde injections from the Allen Mouse Brain Connectivity dataset. We provide CC-MDJ-IO as well as CN-MDJ-IO topographical and density input-output maps, and we show that CC and CN can converge their projections onto the same olivary-projecting MDJ neurons. These data establish the MDJ as an important central hub in wide-spread cerebro-cerebellar communication supporting integration of multiple loops engaging the IO.

## Results

### Projection from MDJ to IO

In mammals, the MDJ is recognized as a dome-shaped area wrapping around the FR at the border of the mesencephalon and diencephalon (Carlton et al., 1982, Onodera, 1984). In the current study on the murine MDJ, we adhere to the terminology in higher mammals such as cats (Onodera and Hicks, 2009, Burman et al., 2000, Strominger et al., 1979, Onodera, 1984), for which the subregions of the MDJ have been more accurately and consistently described than in rats (Brown et al., 1977, Rutherford et al., 1984, Ruigrok, 2004). More specifically, the MDJ in mice forms a continuous rostro-caudal cell column, starting rostrally at the prerubral field (PR) and rostral interstitial nucleus of the medial longitudinal fasciculus (RI) located ventrally to the FR, dwindling medially covering areas of the medial accessory nucleus of Bechterew (NB) and the nucleus of Darkschewitsch (Dk), and ultimately transcending dorsally and caudally on the border of the peri-aquaductal grey (PAG) in the interstitial nucleus of Cajal (InC) and medial accessory oculomotor nucleus (MA3) (Figure 1 - figure supplement 1). Instead, the murine IO is located in the ventral medulla oblongata and consists, just like those of all other mammalian species, of several sheaths of neuropil that form separate subnuclei, including the principal olive (PO), medial accessory olive (MAO), and dorsal accessory olive (DAO) (De Zeeuw et al., 1998).

To uncover the precise topography of the MDJ projection to the IO in mice, we started with retrograde tracing experiments by injecting 1% cholera toxin B (CTB) solution unilaterally in the different olivary subnuclei (n = 7 mice, Figure 1A, B). In addition to some scattered labelling throughout the midbrain, we observed compact, yet overlapping, retrogradely labelled cell populations distributed across various subregions of the ipsilateral MDJ, extending for approximately 1.2 mm in the sagittal plane (Figure 1C, D; Figure 1 - figure supplement 2). In most cases, retrogradely labelled neurons were situated along the rostrocaudal cell column of the PR, RI, NB, Dk, InC and MA3 (Figure 1C, D and Figure 1 - video supplement). The injections that covered the PO received the most prominent projections from the rostroventral subregions of the MDJ, whereas those in the rostral MAO resulted in more retrograde labelling in the more intermediate and caudal subregions of the MDJ (Figure 1 - figure supplement 2). These findings were corroborated by analyses of the outcomes of the injections that were made with the use of different stereotactic coordinates in the rostrocaudal plane.

**Figure 1.**
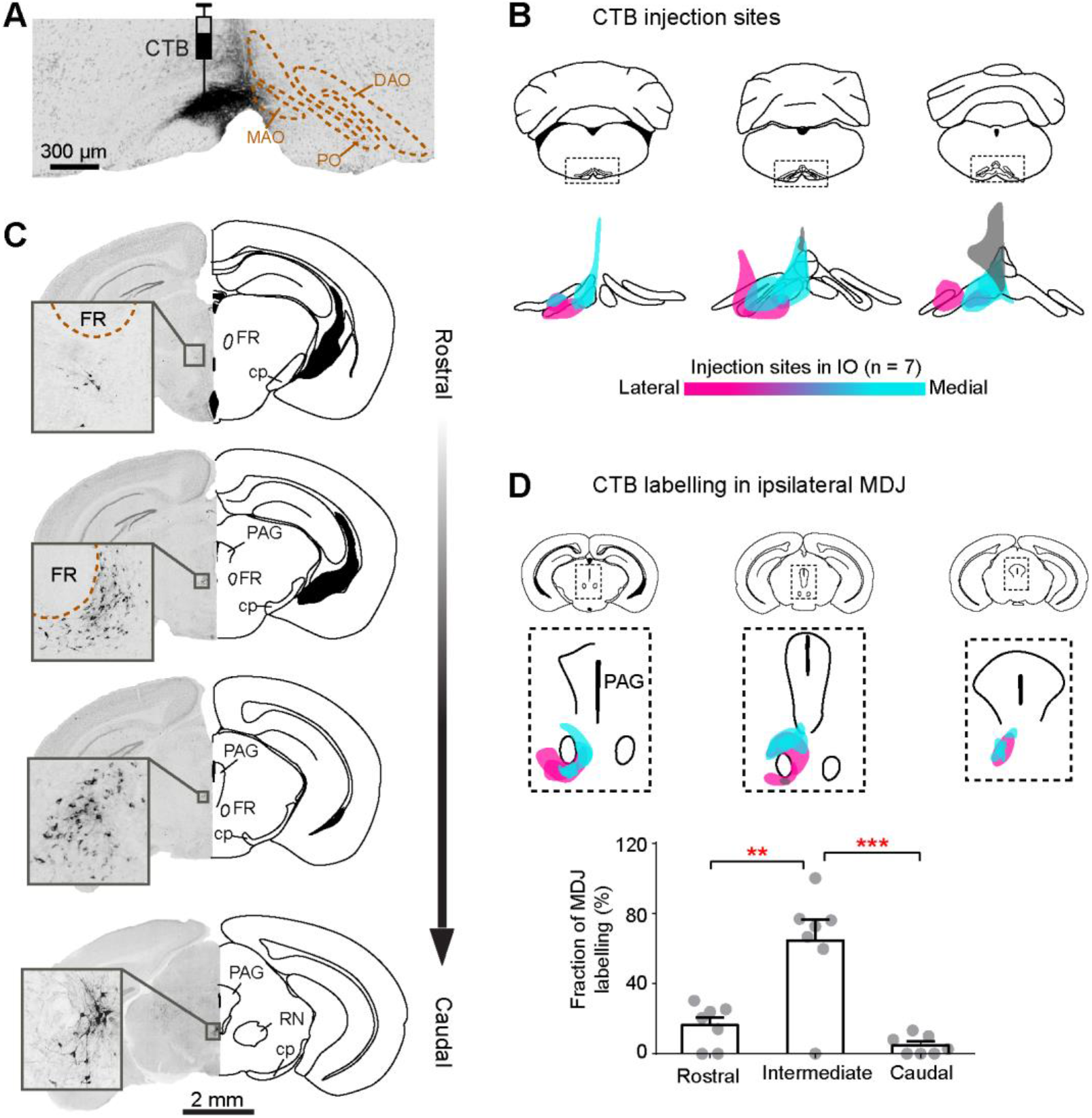
CTB injections in the IO retrogradely label MDJ neurons. (A) Coronal section of IO from a representative mouse, showing CTB injection in the MAO. (B) Summary of the IO injections for seven mice. Magenta, cyan and grey: injections mainly cover PO (n = 2, magenta), rostral MAO (n = 3, cyan) and caudal MAO (n = 2, grey), respectively. (C) Retrogradely ladled neurons in the MDJ at four anteroposterior levels. Sections are obtained from the injection in (A). (D) Summary of CTB labelled neurons in the MDJ regions (upper, overlay of seven mice) and their anteroposterior distribution (lower, mean ± s.e.m.). \*\**P* < 0.01, \*\*\**P* < 0.001, one-way ANOVA, Sidak’s multiple comparisons test.

Notably, no or only very few retrogradely labelled cells were observed within the confines of the magnocellular red nucleus (Figure 1C, D), which is in line with an earlier description in rat (Rutherford et al., 1984, Ruigrok, 2004). In addition to the topographical differences highlighted above, we observed slight differences in the densities of the projections. When we quantified the CTB-labelled cells at distinct rostrocaudal levels of the MDJ in the seven mice, we observed that labelling was most abundant at the intermediate level, comprising the medial accessory nucleus of Bechterew and nucleus of Darkschewitsch (Figure 1D, intermediate versus rostral and caudal level, one-way ANOVA, Sidak’s multiple comparisons test, *P* < 0.05 for both comparisons).

Given that some of our injections diffused slightly into some of the adjacent structures of IO, *i.e.*, the reticular formation of the medulla oblongata, we did additional control experiments in which we injected CTB in the overlying caudal gigantocellular and paramedial reticular nucleus, while avoiding the IO (n = 2 mice, Figure 1 - figure supplement 3A). In these experiments we observed only a very few CTB-labelled cells in the caudal MDJ (Figure 1 - figure supplement 3B), indicating that the MDJ projections described above presumably indeed target virtually exclusively the IO. All together, we conclude that in the mouse a topographic projection from the MDJ to PO and rostral MAO can be recognized, similar to the cat (Onodera, 1984).

### Projection from CC to MDJ

As the projections from the MDJ subregions to the IO subnuclei are organized in a topographical manner (Figure 1), we wanted to find out to what extent the projection from the CC to the MDJ is similarly organised. To this end, we first exploited data gathered for the generation of the Allen Mouse Brain Connectivity Atlas (Allen Atlas in short). This dataset consists of anterograde AAV-GFP viral tracing from various areas of the CC to downstream structures in both wild types (circles) and transgenic mice (squares) (Oh et al., 2014). In our analyses (n = 167 mice), we focused on the cases in which a projection from the CC to the MDJ could be observed (n = 89/167 mice), taking the injection volumes into account (see Materials and Methods for details). We first plotted the CC injections dependent on the rostro-caudal projection of terminals that they provided in the MDJ (Figure 2A), yielding the overall topography on a flattened map of the mouse CC (Figure 2B) (Aoki et al., 2019). Along the 1.2 mm longitudinal axis of the MDJ, the caudal MDJ, comprising InC and MA3, receives most of its projections from relatively rostrolaterally located parts of the ipsilateral CC (e.g., rostral parts of the primary and secondary motor cortices as well as primary and secondary somatosensory cortices; see cyan markers in Figure 2B); whereas axon terminals in the rostral MDJ, covering parts of the PR and RI, originate mainly from injections in either the medial and caudal parts of the ipsilateral CC (e.g., cingulate, retrosplenial and infralimbic cortices as well as specific parts of the auditory and visual cortex; see magenta markers). The intermediate subregions of the MDJ, including NB and DK, also receive input from parts of the primary and secondary motor cortices as well as primary and secondary somatosensory cortices, but those are situated posterior to the analogous sources of the CC that project to the caudal MDJ (see structures in Figure 2B marked with mixed cyan and magenta colours). In addition to all these prominent ipsilateral projections, there were also many contralateral projections. Interestingly, the topography of these contralateral projections was virtually all symmetric to the ipsilateral ones, highlighting the relevance of bilateral control (Figure 2B, right panel).

**Figure 2.**
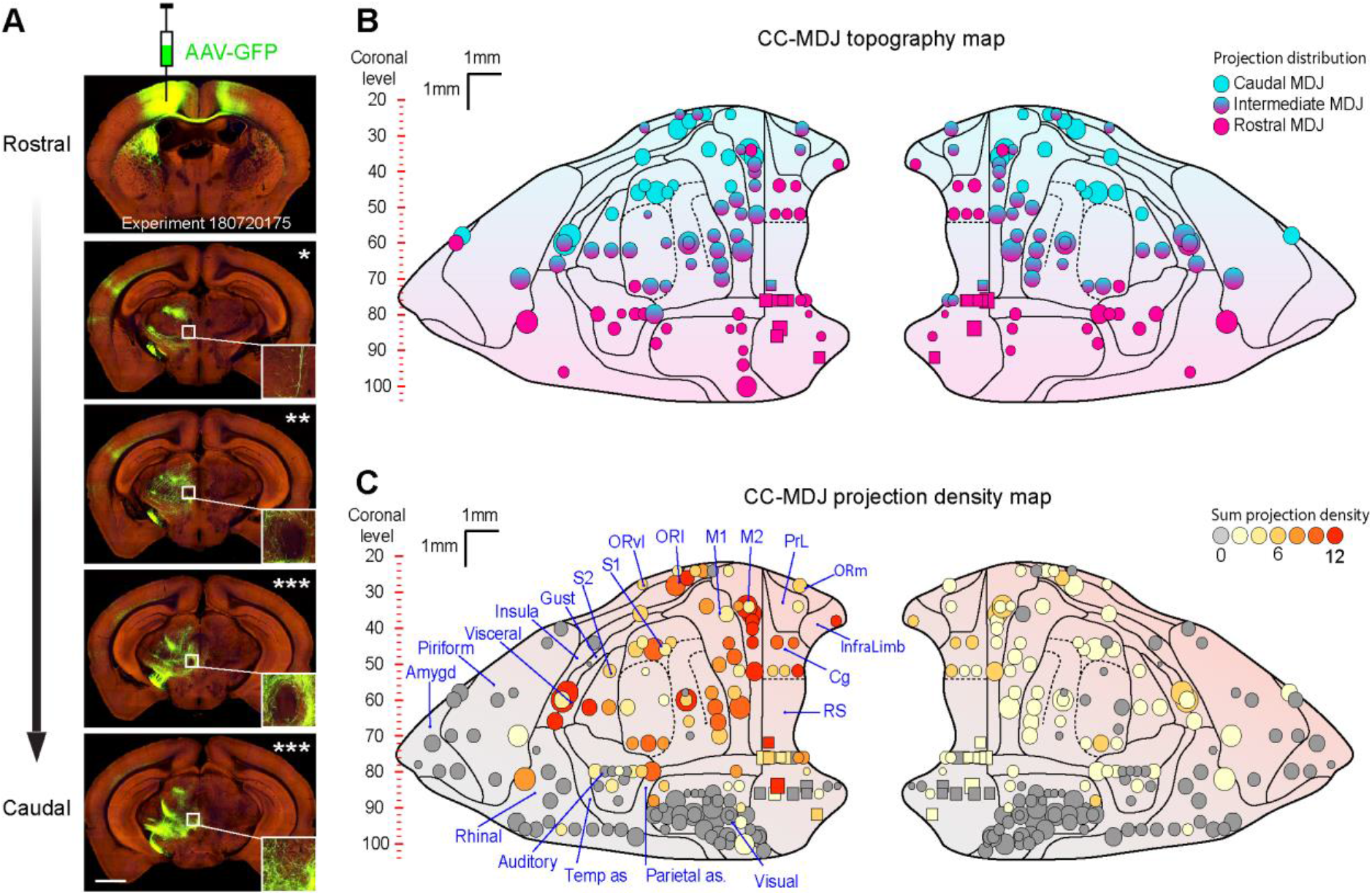
Viral anterograde tracing demonstrates CC-MDJ topographic and density organisations. (A) Images of an example case from the Allen Mouse Brain Atlas with an injection in the motor cortex (top) and projections at anteroposterior levels of the MDJ. The number of asterisks represents cortical axon density in the MDJ (insert, see quantification in the Materials and Methods). (B) CC-MDJ topography map of cortical injections on our standard flattened CC representation (Aoki et al., 2019). Each marker represents one injection in the CC (wild types in circle, n = 6/167; transgenics in squares, n = 83/167); marker size indicates injection volume (see Materials and Method); colour coding suggests axonal projection in the anteroposterior MDJ. (C) CC-MDJ density map of cortical areas projecting onto the MDJ. The different density levels are represented by the colour of the circles and squares on the standard flattened scheme of the CC. Note that the cases exhibiting no MDJ projection (grey markers, n = 78/167) were not illustrated in panel (B). Abbreviations: Amygd: amygdala, Auditory: auditory cortex, Cg: cingulate cortex, Gust: gustatory cortex, InfraLimb: infralimbic cortex, Insula: insular cortex, M1: primary motor cortex, M2: secondary motor cortex, ORl: lateral orbital cortex, ORm: medial orbital cortex, ORvl: ventrolateral orbital cortex, Parietal as: parietal association cortex, Piriform: piriform cortex, PrL: prelimbic cortex, Rhinal: rhinal cortex, RS: retrosplenial cortex, S1: primary somatosensory cortex, S2: secondary somatosensory cortex, Temp as: temporal association cortex, Visceral: visceral cortex, Visual: visual cortex.

To estimate the relative prominence of the MDJ projections from the different cortical regions, we presented the density of these projections of the Allen Atlas with colour coding onto the same map. The most densely labelled terminal fields covering the MDJ resulted from mice with injections in the ipsilateral infralimbic, cingulate, secondary motor, sensory, lateral orbital, gustatory and/or visceral cortical areas, but the injections in many of the other areas also resulted in some labelling of efferent axons located in the ipsilateral MDJ (n = 89 mice, coloured markers in Figure 2C). In contrast, many injections in various parts of the CC resulted in no labelling of the MDJ; these included for example specific parts of the piriform, visual, auditory, and parts of the somatosensory cortices (n = 78 mice, grey markers in Figure 2C). Finally, the symmetry between the ipsilateral and contralateral projections that was revealed in the topography map (Figure 2B) was to some extent also present in the respective density maps, with the contralateral side showing a consistently lower number of efferent fibres (Figure 2C). Hence, by evaluating a large number of cases our retrieval analyses yielded the CC-MDJ projection maps, establishing their topographic relations and density ‘hot spots’.

Next, we wanted to verify the anterograde approach of the Allen Atlas with retrograde tracing experiments. To this end, we performed small unilateral CTB injections (n = 9) targeting the MDJ region at the various caudo-rostral levels (Figure 3A) and examined the retrograde labelling in the CC (Figure 3B). In addition to some retrogradely labelled cells in the lateral habenular nucleus, precommissural nucleus, parafascicular thalamic nucleus, and posterior paraventricular thalamic nucleus, CTB labelled cells were observed in various CC regions predominantly, but not exclusively, on the ipsilateral side (Figure 3B). In terms of distribution density, cortical labelling was in general most abundant in the motor, cingulate, primary somatosensory, insula, prelimbic and infralimbic cortices, labelling more than 5% of the total CTB-labelled neurons in the CC (Figure 3B). The injections at the caudal and rostral level of the MDJ showed relatively high numbers of retrogradely labelled cells in the rostrolateral and medial areas of the CC, respectively (compare Figure 3C, D with Figure 3E, F, respectively). For example, the CTB injections in the caudal and rostral MDJ provided particularly robust retrograde labelling in parts of the motor and cingulate cortices. These results are in line with the topography and density maps of the CC-MDJ projection revealed with anterograde tracing (Figure 2).

**Figure 3.**
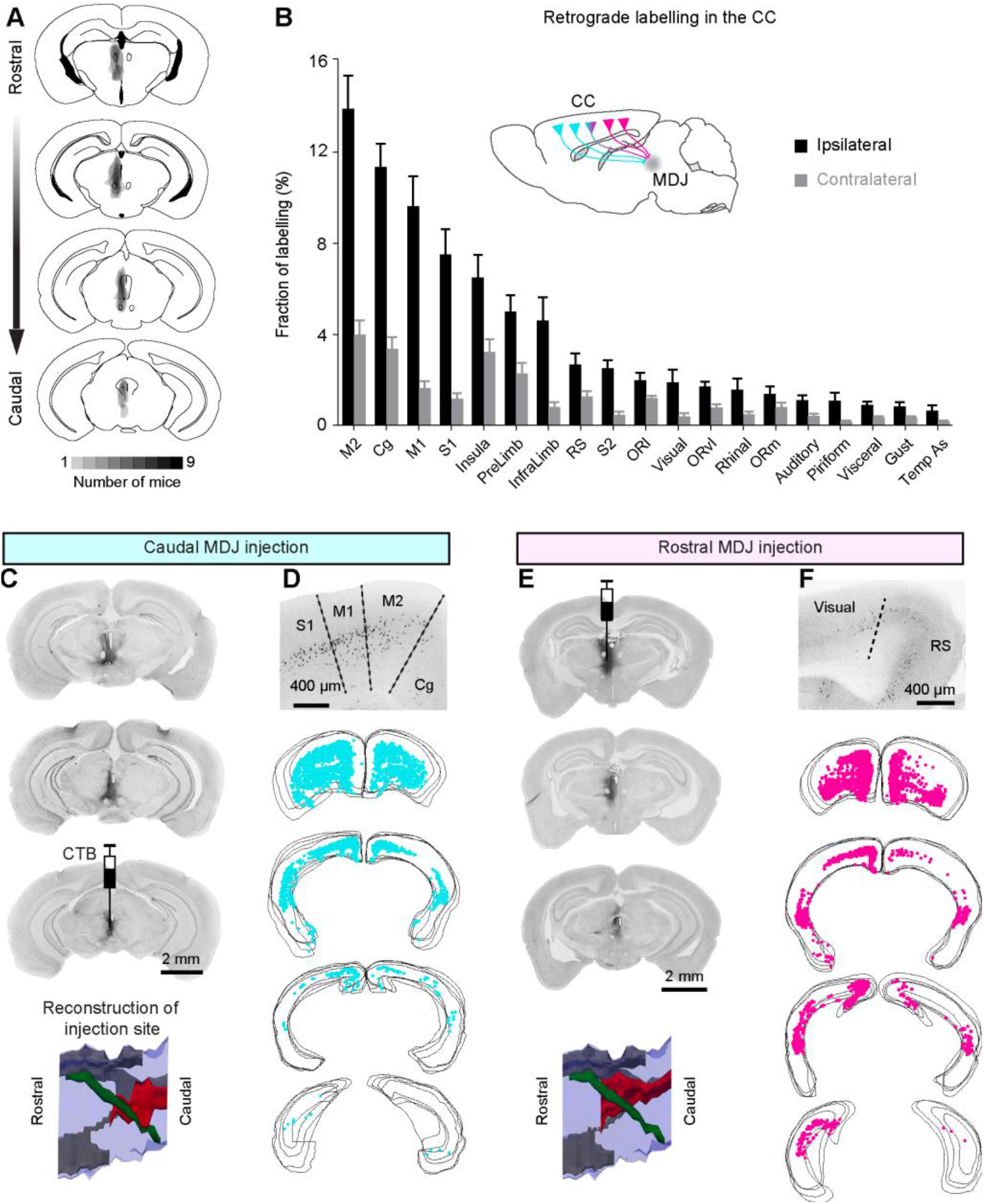
CTB retrograde tracing from the MDJ to the CC. (A) Summary of CTB injections in the MDJ regions (n = 9 mice). (B) Quantification of retrogradely labelled cells in the CC (mean ± s.e.m.). Light grey and dark grey represent fractions of total labelled cells on the contralateral and ipsilateral sides, respectively. (C-F) Comparison of retrograde cortical labelling from a caudal MDJ-injected case and a rostral MDJ-injected case. In the lateral view of injection site reconstruction (bottom, C and E), red: CTB injection site, green: FR, grey: third ventricle, blue: midbrain contour. Example cortical sections (D and F) show CTB labelling ipsilateral to the injection sites. In the serial plotting of cortical labelling (D and F), each panel include four consecutive sections. See abbreviations in **Figure 4**.

Given that some of the CTB injections also included the parafascicular thalamic nucleus and PAG, which are known to receive afferent projections from the cingulate, prelimbic and infralimbic cortices (Royce et al., 1991, Teune et al., 2000, Gonzalo-Ruiz et al., 1990, Cornwall and Phillipson, 1988, Mandelbaum et al., 2019, An et al., 1998), we did additional control experiments by targeting these nuclei selectively without hitting the MDJ (Figure 3 - figure supplement 1A). In principle, these nuclei might also constitute an intermediary hub between the CC and the IO. In these control injections surrounding the MDJ, we did indeed observe abundant retrograde labelling in the cingulate and limbic cortices as well as in the medial prefrontal cortex, but we hardly observed any axonal labelling in the IO (Figure 3 - figure supplement 1B, C), excluding the possibility that the parafascicular thalamic nucleus and adjacent PAG form major hubs between the CC and IO. Taken together with the anterograde experiments from the CC described above (Figure 2), our retrograde tracing experiments highlight that the projection from the CC to the MDJ concerns very large, but not all, parts of the CC and that this input is topographically organised and dense. At the same time, it should be noted that when elucidating the connectivity of the CC with the IO via the MDJ, it is important to take a double experimental approach, exploiting both anterograde and retrograde tracing methods to avoid potential caveats.

### Topographic connectivity between CC, MDJ and IO

To be able to better compare the topography of the inputs from the CC to the different MDJ regions with that of the MDJ outputs to the IO, we next plotted the anterograde labelling in the different olivary subdivisions (Figure 4). The CTB injections at the rostral and caudal levels of the MDJ showed relatively high numbers of anterogradely labelled fibres in the ipsilateral PO and rostral MAO, respectively (compare e.g., Figure 4A, B). Interestingly, the fibres that traversed from the rostral MDJ to the PO appeared to be routed via the central tegmental tract, whereas those of the caudal MDJ traveling towards the rostral MAO were predominantly located in the medial tegmental tract, which is in line with earlier descriptions in higher mammals (Burman et al., 2000, Ruigrok, 2004). When putting the data provided by the injections in the IO (Figure 1), the CC (Figure 2), and MDJ (Figures 3 and 4), together, the picture emerges that the rostromedial and caudal parts of the CC predominantly project to the PO via the PR and RI, that the rostrolateral parts of the CC predominantly project to the rostral MAO via the InC and MA3, and that the in-between parts of the CC predominantly project to overlapping parts of the PO and MAO via the BN and DK (Figure 4E).

**Figure 4.**
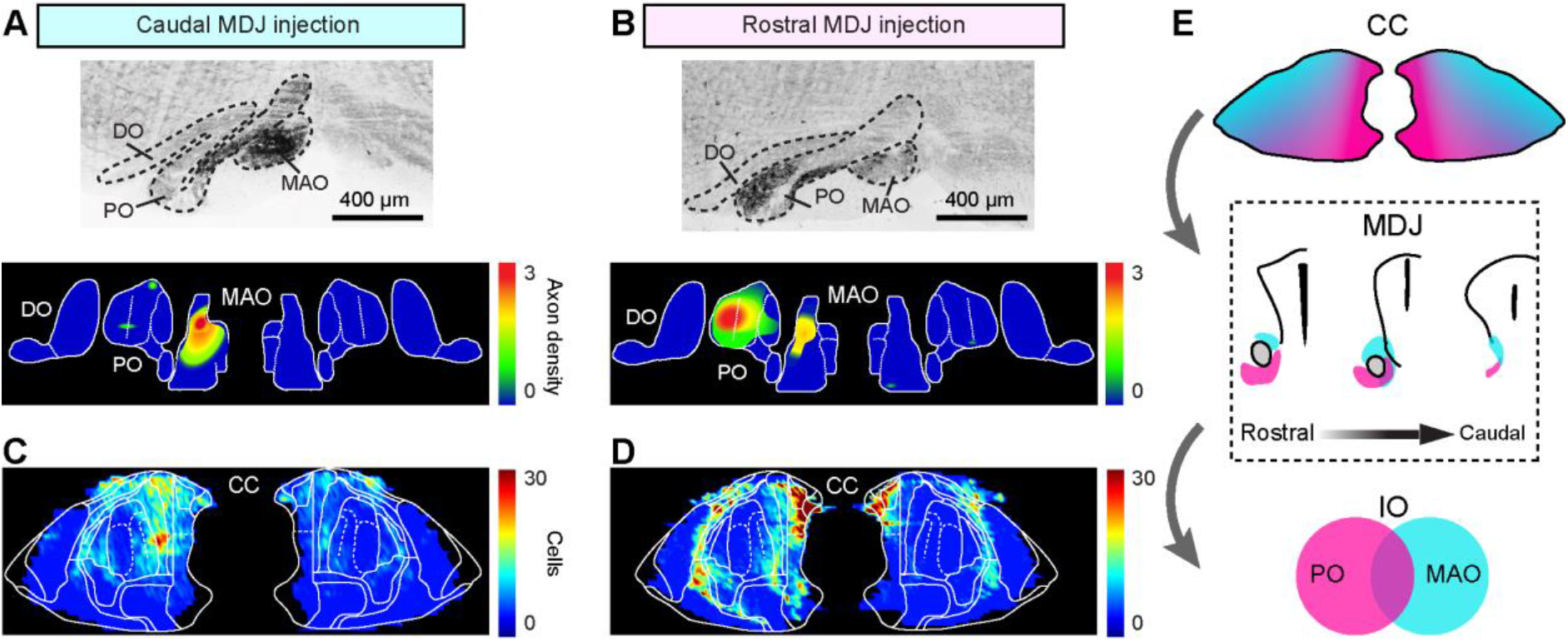
CC-MDJ-IO topography by injecting CTB in anteroposterior MDJ. (A and B) Example sections (upper) and density maps (lower) of anterogradely labelled MDJ axons in the IO. (C and D) Density maps of retrogradely labelled neurons in the CC from the caudal and rostral MDJ injections. (E) Schematic of CC-MDJ-IO topographic organisation. Abbreviations: DO: dorsal accessory olive, MAO: medial accessory olive, PO: principal olive.

### Projection to MDJ from CN

The CTB injections in the MDJ described above also provided substantial retrograde labelling in the contralateral CN (Figure 5A-C). More specifically, the rostral and caudal MDJ injections resulted in retrogradely labelled cells in predominantly the contralateral dentate nucleus (DN) and posterior interposed nucleus (PIN), respectively (Figure 5B, C). Each of them resulted in more than 25% of the total CN labelling (Figure 5A). Approximately 15% of the total CN labelling was found in the anterior interposed nucleus (AIN), including the dorsolateral hump (Figure 5A). Notably, the variability of retrograde labelling was relatively high in the fastigial nucleus (FN) (s.e.m. in FN = 6.2 versus 3.1, 3.3 and 2.4 in the PIN, AIN and DN, respectively). This may partly reflect the fact that two of the nine injections in the MDJ regions diffused into the oculomotor area and PAG, which are known to receive abundant FN input (Gonzalo-Ruiz et al., 1990, Teune et al., 2000, Fujita et al., 2020); without these two animals only 15.1 ± 1.9% of the FN cells were labelled.

**Figure 5.**
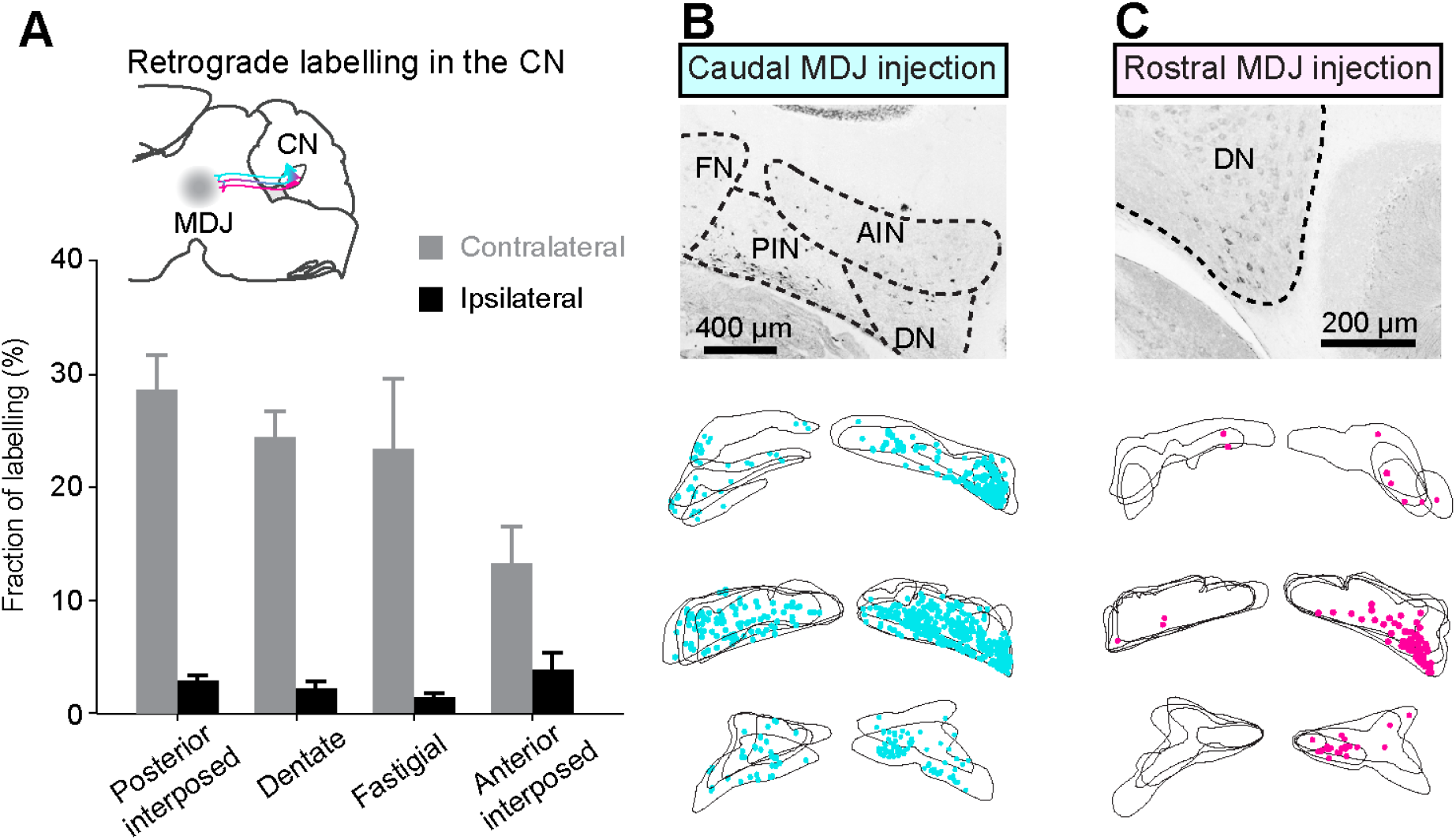
Retrograde CTB labelling from the MDJ to the CN. (A) Summary of CTB labelled neurons in the CN (mean ± s.e.m., n = 9 mice). (B and C) Comparison of retrograde CN labelling from a caudal MDJ-injected case and a rostral MDJ-injected case. Example sections show labelling in the contralateral CN. In the serial plotting of CN labelling, each panel include four consecutive sections.

### CC and CN projections converge onto MDJ neurons projecting to IO

Our CTB injections in the MDJ simultaneously provided retrograde labelling of cells in the CC and CN as well as anterograde labelling of axonal fibres in the IO (Figures 3–5). This raises the possibility that IO-projecting neurons in the MDJ may function as a central hub in the brainstem, onto which both CC and CN converge. To further clarify the relationship between MDJ input and output, we used a gene-modified transneuronal rabies strategy (Wall et al., 2010, Kim et al., 2016, Callaway and Luo, 2015), in which we retrogradely traced neurons in the CC and CN that send axon terminals to MDJ cells projecting to the IO (n = 4 mice). We first injected Cre-dependent helper virus AAV8-CAG-FLEX-TCB, AAV8-CAG-FLEX-oG in the MDJ and hSyn-AAVretro-Cre-eBFP in the ipsilateral IO, allowing MDJ neurons that project to the IO to express both rabies glycoprotein (oG) and avian acceptors (TVA) (Figure 6A). Next, following four weeks of incubation, we injected glycoprotein gene-deleted rabies pseudotyped with the avian sarcoma leucosis virus glycoprotein EnvA (RV-CMV-EnvA-ΔG-eGFP) in the MDJ to induce cell-specific, retrograde transneuronal labelling (Figure 6A). Indeed, after another week of incubation, rabies transfected a group of MDJ neurons (starter cells) that expressed TVA on their membranes (Figure 6B). The location of these starters cells was in accordance with that of the IO-projecting MDJ neurons (Figure 1), i.e., a cell column surrounding the FR (Figure 6B). Transsynaptic retrograde rabies labelling was found in a variety of regions located at different distances. At short-distance, we observed labelling in the zona incerta, the prerubral field and the posterior commissure nuclei (Figure 6B), while at longer distances, we found labelling in both the CC and CN (Figure 6C and D). The retrogradely labelled cells in the CC and CN were located predominantly on the ipsilateral and contralateral side to the MDJ, respectively. In the CC, these comprised only layer-V pyramidal neurons with their characteristic angular-shaped somata, multipolar dendritic trees and elongated axons (Figure 6C); these neurons were distributed in specific parts of multiple cortical regions, including for example the sensory, motor, cingulate and retrosplenial cortices (Figure 6C). In the CN, these comprised the larger, presumably excitatory, neurons in the DN and PIN (Figure 6D). Thus, in line with the analyses of the Allen Atlas data and the CTB experiments described above (Figures 2–5), the transneuronal rabies-tracing experiments demonstrated the existence of IO-projecting neurons in the MDJ that receive direct input from the CC and also the existence of IO-projecting neurons in the MDJ that receive direct input from the CN.

**Figure 6.**
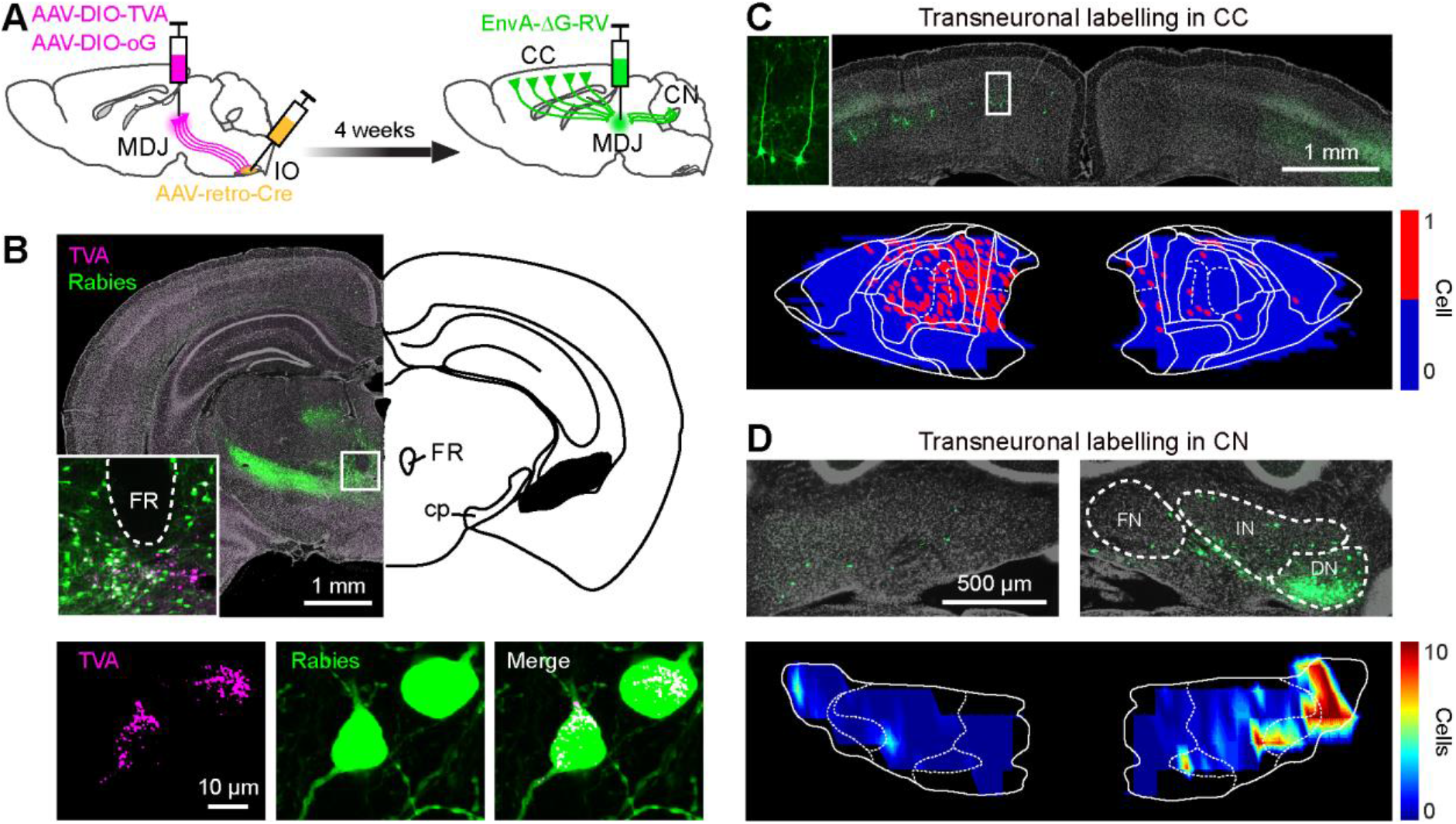
Monosynaptic rabies tracing revealing CC and CN inputs on the olivary-projecting MDJ neurons. (A) Schematic of transneuronal viral tracing strategy (n = 4 mice). (B) Coronal sections of a representative mouse at the midbrain level (upper, insert suggesting MDJ), showing rabies primary labelling (lower, TVA and rabies co-labelled) and local secondary labelling. (C) Coronal section (upper) and plotting map (lower) of transneuronal rabies labelling in pyramidal cells (insert) of motor cortex. (D) Same as (C), but for labelling in the CN. DN: dentate nucleus, FN: fastigial nucleus, IN: interposed nucleus.

The experiments described above provided supportive evidence for the MDJ to operate as a CC-CN converging hub, but they did not allow us to unequivocally address whether individual IO-projecting neurons in the MDJ can receive input from both, the CC and CN. To explore whether CC and CN axons can converge onto the same olivary-projecting MDJ neurons, we performed triple viral-tracing experiments (n = 3 mice) combining retrograde tracing of AAVretro-GFP from the IO with anterograde tracing of AAV1-CAG- GFP_sm_-myc and AAV1-CB7-RFP in the CC (motor cortex) and CN (DN and PIN), respectively (Figure 7A). AAVretro-GFP virus was taken up by MDJ axons in the IO and subsequently transported retrogradely to the somata of neurons in MDJ areas that also received input from anterogradely labelled axons derived from the CC and/or CN (Figure 7B). Even though the inputs from the CC and CN to the most caudal tip of the MDJ were partly segregated, in the more rostral and intermediate subregions of the MDJ there was a prominent convergence of CC and CN inputs to olivary projecting neurons (see e.g., 2nd panel on the left of Figure 7B with prominent triple labelling in NB). This conclusion was supported by quantification of the fractions of IO-projecting MDJ cells that received input from CC or CN axons at the different rostrocaudal levels (Figure 7C). Moreover, we found the same topographic distribution of terminals derived from the various CC areas and CN areas as found following AAV-GFP and CTB injections (Figure 2 and Figure 5, respectively). It is hard to exclude that some of the light microscopic profiles in the MDJ were false-positively interpreted as axon terminals (instead of passing fibres), but we are rather confident that most of the CC versus CN identifications were correct because of their peculiar postsynaptic distribution and size. Indeed, whereas CC boutons were mostly apposed to secondary dendrites, CN axon terminals preferably terminated on the somata and primary dendrites of MDJ neurons (Figure 7D). Likewise, the average size of CC profiles terminating on IO-projecting neurons in the MDJ (1.07 ± 0.15 μm^2^, n = 27) was significantly smaller (two-sample *t-test*, *P* = 0.0005) than that of CN profiles (2.22 ± 0.24 μm^2^, n = 37) (Figure 7E), which is in line with that of previous electron microscopic studies (De Zeeuw and Ruigrok, 1994). Taken together, our triple-tracing experiments indicate that axon terminals originating from the CC and the CN can converge onto the same individual IO-projecting neurons in the MDJ, yet with differential distribution patterns at the subcellular level.

**Figure 7.**
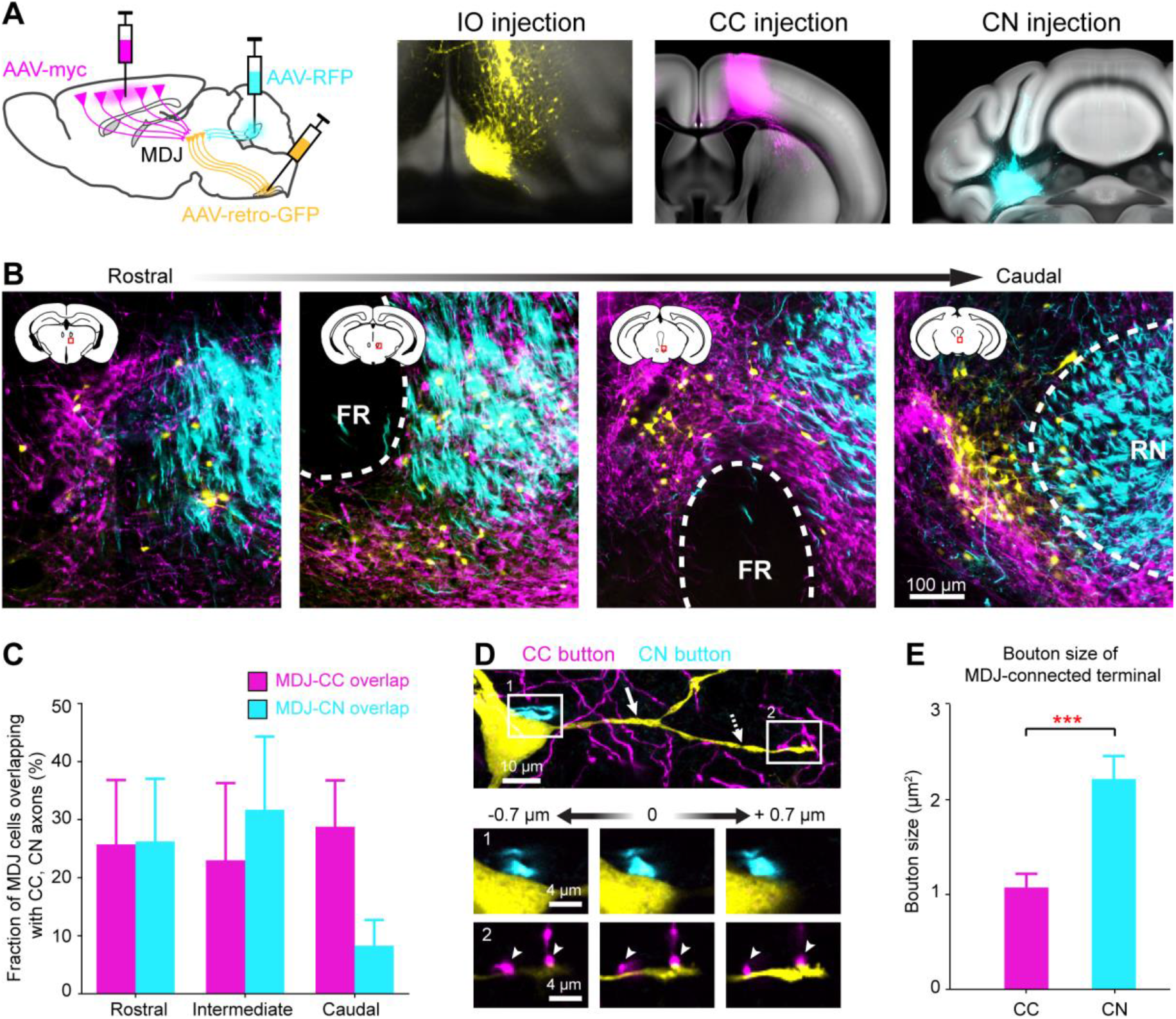
Convergence of CC and CN terminations on the olivary-projecting MDJ neurons. (A) Schematic of tracing strategy. Coronal sections (registered in the Allen CCF) showing injections of retrograde AAV-GFP in the IO, anterograde AAV-myc in the motor cortex and AAV-RFP in the cerebellar interposed-dentate complex. (B) Coronal sections of MDJ regions from an example mouse showing retrogradely-labelled cells (olivary-projecting cells, yellow) distributing in the CC (magenta) and CN (cyan) axons. (C) Colocalization quantification of olivary-projecting cells and CC, CN axons at anteroposterior MDJ levels (n = 3 mice, mean ± s.e.m.). (D) Confocal images of an olivary-projecting MDJ neuron from (B) showing close synaptic contact with CN axon on its soma (insert 1) and with CC axons on its secondary dendrite (insert 2). Solid arrow: primary dendrite, dashed-line arrow: secondary dendrite. (E) Size comparison of CC and CN buttons exhibiting close contacts with IO-projection MDJ cells (n = 27 for CC buttons, n = 37 for CN buttons, mean ± s.e.m., two-sample *t-test*, \*\*\**P* < 0.001).

## Discussion

When considering how the CC may influence cerebellar activity, the first major hub that comes to mind is the pons, which is the main source of mossy fibres for the cerebellum (Henschke and Pakan, 2020, Coffman et al., 2011, Ruigrok, 2004). The pons receives prominent topographic projections from a wide variety of cortical regions (Leergaard, 2003, Leergaard et al., 2000, Wiesendanger and Wiesendanger, 1982) and has been shown to exert a powerful control over the lateral hemispheres of the cerebellar cortex and dentate nucleus during various types of behaviours, including planning and coordination of limb and eye movements (Guo et al., 2020, Chabrol et al., 2019, Ohmae et al., 2017). Here, we show unique topographic projections from the CC to the cerebellar cortex via the MDJ and IO, providing the climbing fibres to the Purkinje cells in the cerebellum. Exploiting advancements in transneuronal viral tracing approaches as well as emerging large-scale databases on brain connectivity (Allen Atlas), we have been able to elucidate the topography of both the CC-MDJ-IO loop and the CN-MDJ-IO loop. Our data obtained in mice expands on existing data on the individual connections between CC and MDJ, CN and MDJ, as well as between MDJ and IO in a wide variety of species. More specifically, we demonstrate that rostromedial and caudal parts of the CC project predominantly to ventrorostral parts of the MDJ, which in turn provide a dense projection to the PO, whereas the more rostrolateral parts of the CC project predominantly to the caudodorsal parts of the MDJ, which in turn target predominantly the rostral MAO. The topography of this CC-MDJ-IO loop aligns well with that of the CN-MDJ-IO loop (De Zeeuw and Ruigrok, 1994). Whereas the dentate nucleus receiving climbing fibre collaterals from the PO projects most prominently to the rostral MDJ, the posterior interposed nucleus, which receives its collaterals from the rostral MAO, projects most prominently to the caudal MDJ. Together, these findings highlight that the integration of cerebral cortical with cerebellar information is not only mediated by a loop via the pontine mossy fibre system, but also by MDJ loops engaging the olivary climbing fibre system.

### Topography of CC-MDJ-IO loop

By systematically analyzing a large number of cases with cortical anterograde viral injections from the Allen Mouse Brain dataset, and verifying this by performing retrograde CTB in the MDJ, we identified the topography of the cortical regions that provide input to the MDJ. In general, the rostromedial and caudal parts of the CC predominantly project to the rostroventral areas of the MDJ and the more rostrolateral parts of the CC prominently project to the caudodorsal areas of the MDJ. Our CTB injections in the rostral and caudal aspects of the MDJ in mice are in line with earlier results obtained in monkey (Leichnetz et al., 1984), cat (Nakamura et al., 1983b, Saint-Cyr, 1987, Rutherford et al., 1989, Onodera and Hicks, 1996), rat (Murphy and Deutch, 2018, Hoover and Vertes, 2007), as well as mice (Kubo et al., 2018, Oh et al., 2014), as the most prominently distribution of retrogradely labelled cells were found in the somatosensory, motor and premotor cortices as well as the cingulate and prefrontal areas.

Likewise, the topographic connections we showed here between the MDJ and IO for mice have also been studied in different species such as monkey (Strominger et al., 1979, Onodera and Hicks, 2009, Burman et al., 2000), cat (Mabuchi and Kusama, 1970, Saint-Cyr and Courville, 1981, Saint-Cyr, 1987, Onodera, 1984) and rat (Carlton et al., 1982, Rutherford et al., 1984, Voogd and Ruigrok, 2004). In cat, retrograde tracing studies from selective areas of the IO complex have demonstrated subpopulations of labelled neurons in the MDJ area, distributed over a wide variety of nuclei, including the parvocellular red nucleus, medial accessory nucleus of Bechterew, nucleus of Darkschewitsch, suprarubal reticular formation, nucleus of the fields of Forel and interstitial nucleus of Cajal (Onodera, 1984, Saint-Cyr and Courville, 1981, Walberg and Nordby, 1981, Condé and Condé, 1982, Onodera and Hicks, 2009). For example, areas like the medial accessory nucleus of Bechterew and the nucleus of Darkschewitsch project prominently to the PO and rostral MAO, respectively (Onodera, 1984). Studies in human (Burman et al., 2000, Strominger et al., 1979, Massion, 1967, Onodera and Hicks, 2009) and non-human primates (Onodera and Hicks, 2009, Burman et al., 2000, Strominger et al., 1979) reported results that are largely comparable to those in the cat. Interestingly, in primates the tract that carries the fibres from the parvocellular red nucleus projecting to the PO, the central tegmental tract, has expanded proportionally, standing out against the medial tegmental tract that holds the fibres traversing from the nucleus of Darkschewitsch to the rostral MAO (Burman et al., 2000, Voogd and Ruigrok, 2004). Here, in mice, we find a similar location of the fibres traversing from the rostral MDJ to the PO, next to those traversing from the caudal MDJ to the rostral MAO, but this route does not show the evolutionary expansion of the central tegmental tract seen in higher mammals. In our view, the prominent topography in the connectivity between CC, MDJ and IO as well as the expansion of the parvocellular red nucleus and the central tegmental tract in higher mammals underscores the emerging relevance of the MDJ as an intermediary hub in cerebrocerebellar processing.

### Topography of CN-MDJ-IO loop

Our tracing experiments showed that the olive-projecting cells in the MDJ receive not only topographically organized inputs from different areas of the CC, but also inputs from the CN in an analogous way. In general, our findings in mouse are consistent with previous tracing results in rat (Teune et al., 2000, Bentivoglio and Kuypers, 1982, Berretta et al., 1993, Ruigrok and Teune, 2014), cat (Fukushima et al., 1986, De Zeeuw and Ruigrok, 1994), and monkey (Stanton, 1980, Gonzalo-Ruiz et al., 1988, Kalil, 1981), showing that the cerebellar projections onto the MDJ nuclei are also topographically organized. The cells in the rostral MDJ that get input from rostromedial and caudal parts of the CC receive inputs from predominantly the dentate nucleus, whereas the neurons in the caudal MDJ that are more prominently connected with rostrolateral parts of the CC mainly receive input from the interposed nucleus. These findings on the topography were corroborated by the density studies following CTB injections in the MDJ, in which we found that the amount of retrogradely labelled cells in the CN was positively correlated with the amount of anterogradely labelled terminals in the IO. Moreover, the reciprocal connections between the different CN and olivary subnuclei are also in line with the topography described above in that the dentate nucleus is mainly connected with the ventral and dorsal lamellae of the PO (Apps and Hawkes, 2009), which receives input from the rostral MDJ, and that the interposed nucleus is mainly connected with the rostral MAO (Apps and Hawkes, 2009), which receives input from the caudal MDJ. Thus, the two loops, i.e., the CC-MDJ-IO loop and the CN-MDJ-IO loop, are connected with each other in a topographically consistent way.

### Convergence of cerebral and cerebellar input to the MDJ

In addition to elucidating the topography of the CC-MDJ-IO and CN-MDJ-IO loops, we also show here for the first time that these two loops share at least in part the same MDJ neurons; indeed, CC pyramidal cells and contralateral CN projection cells converge upon the same MDJ neurons that project to the IO. This conclusion was based on three lines of evidence. First, our small CTB injections in the MDJ simultaneously provided retrograde labelling of cells in the CC and CN as well as anterograde labelling of axonal fibres in the IO (Figure 4); second, our gene-modified transneuronal rabies approach demonstrated the existence of IO-projecting neurons in the MDJ that do receive direct input from the CC as well as the existence of IO-projecting neurons in the MDJ that do receive direct input from the CN (Figure 6); and third, our triple-tracing experiments (Figure 7) directly revealed individually retrogradely labelled MDJ neurons that received anterogradely labelled axonal input from both the CC and CN, with separately identifiable markers for both inputs. Thereby, our results in mice expand upon classical tracing studies, in which some of the components of the MDJ trajectories were studied in isolation in monkey, cat or rat (de Zeeuw et al., 1989, Leichnetz et al., 1984, Mabuchi and Kusama, 1970, McCrea and Baker, 1985, Nakamura et al., 1983b, Nakamura et al., 1983a, Saint-Cyr, 1987, De Zeeuw and Ruigrok, 1994).

### Role of MDJ and IO in cerebro-cerebellar communication

Multiple lines of research have implicated involvement of the MDJ in CC- and/or cerebellum-dependent behaviors (Halmagyi et al., 1994, Shiraishi and Nakao, 1994, Peschanski and Mantyh, 1983, Wiest et al., 1996). Previous physiological studies provide evidence for direct activation of the IO or increased complex spike activity of Purkinje cells following electrical stimulation of CC regions in awake or anesthetized animals (Kato et al., 1988, Watson et al., 2009, Crill, 1970, Pardoe et al., 2004, Sasaki et al., 1975). Indeed, complex spike activity of Purkinje cells in crus I, crus II, and vermal lobule VII can be facilitated by stimulating the prefrontal (Watson et al., 2009), motor (Ackerley et al., 2006), and/or somatosensory cortices (Kubo et al., 2018), in line with the sources in the CC that provide axonal projections onto the olivary-projecting neurons in the MDJ (see e.g., Figures 2, 6 and 7).

What might be the function of the CC-MDJ-IO loop? The climbing fibres arising from the CC-MDJ-IO loop may provide well-timed, teaching, reward, and/or prediction signals to the Purkinje cells that can be used for complex motor and/or cognitive learning (Heffley et al., 2018, Kostadinov et al., 2019, Ohmae and Medina, 2015). Indeed, an increase of such signals may be used to enhance acquired motor or cognitive responses that are mediated by downbound modules, whereas a decrease of them may be used to drive learning mediated by upbound modules (De Zeeuw, 2021). The increases in climbing fibre activity and associated complex spike activity in the downbound modules may decrease the simple spike activity of Purkinje cells that can drive for example a conditioned, accelerated (Ohmae and Medina, 2015, ten Brinke et al., 2015) or delayed (Chabrol et al., 2019, Gao et al., 2018), motor or cognitive response. Conversely, decreases in climbing fibre activity and associated complex spike activity in the upbound modules can increase the simple spike activity of Purkinje cells that can modulate ongoing motor and/or cortical responses (Voges et al., 2017, Romano et al., 2018). It has been hypothesized that the net-polarity of neurotransmissions downstream of the Purkinje cells in downbound and upbound modules are excitatory and inhibitory, respectively, together facilitating bidirectional control (De Zeeuw, 2021). To what extent the modules of the CC-MDJ-IO loop are implicated in downbound and/or upbound modules remains to be elucidated, but given the mixed upbound-downbound biochemical nature of the D-zones of Purkinje cells in the cerebellar cortex, which receive their climbing fibres from the PO, and the upbound biochemical nature of the C2-zone, which receives its climbing fibres from the rostral MAO, bidirectional control of complex motor responses and/or cognitive responses should indeed be possible (Fujita et al., 2020, Barmack, 2006, Braak et al., 2003, Larson et al., 1969, De Zeeuw, 2021).

The CN-MDJ-IO loop may well support the CC-MDJ-IO loop in serving these timing functions during bidirectional control. Whereas the MDJ provides a robust excitatory input to the electrotonically coupled dendritic spines of the PO and rostral MAO, the CN provides an equally robust inhibitory input to the very same dendritic spines (Ruigrok and Voogd, 1995, De Zeeuw et al., 1998, de Zeeuw et al., 1989). This configuration of a combined excitatory and inhibitory input to excitable spines within a glomerulus renders the activation of olivary neurons particularly sensitive to the timing between these two inputs (De Zeeuw et al., 1998, Negrello et al., 2019). Moreover, as we show in the current study the CN apparently also provides a prominent projection to MDJ neurons that is topographically aligned with their input from the CC. Thus, both at the level of cyto-architecture of the olivary neuropil and at the level of the network connectivity of the CN-MDJ-IO and CC-MDJ-IO loops, the climbing fibre system appears well designed to facilitate precise timing of neuronal activity required for complex functions mediated by cerebro-cerebellar control. It will be interesting to elucidate to what extent processing in the cortico-ponto-cerebellar mossy fibre system lines up with that of the cortico-MDJ-IO-cerebellar climbing fibre system in regulating these functions.

## Materials and Methods

### Animals and viral vectors

Both male and female C57BL/6J mice were used in this study. Animals were 10-20 weeks old and were housed individually in a 12-hour light-dark cycle with food and water *ad libitum*. All experiments were approved by the institutional animal welfare committee of Erasmus MC (15-273-146 and 15-273-147) in accordance with Central Authority for Scientific Procedures on Animals Guidelines.

Adeno-associated virus AAV8-CAG-FLEX-TCB, AAV8-CAG-FLEX-oG and rabies virus RV-CMV-EnvA-ΔG-eGFP were obtained from Salk vector core. AAV1-CAG-GFP_sm_-myc, AAV1-CB7-RFP, AAVretro-CAG-GFP and AAVretro- hSyn-Cre-eBFP were obtained from Addgene.

### Surgical procedures

Animals were anesthetized with 5% isoflurane for induction and 2.5% for maintenance and were fixed on a stereotaxic surgical plate (David Kopf Instruments). Body temperature was kept at 37 ± 0.5 °C constantly during operation. DuraTears (Alcon Laboratories, Inc.) was used to moisture the eyes, and lidocaine (2.5 mg/mL) was locally applied on the skin after removing hair over the scalp. A small vertical incision was applied on the scalp to expose the skull. Animal head was positioned so that the bregma and lambda were well levelled. Stereotaxic coordinates (see Supplementary Table 1) were measured for different injecting targets, then a small craniotomy (Φ = 0.7 mm) was made on the skull. We gently lowered a glass capillary (Φ = 8 μm opening) in the targeted regions and slowly injected about 30 nl cholera toxin B subunit (CTB, 1%, Sigma-Aldrich) or virus. Capillaries were left on the injection sites for approximately 2 mins before being removed from the brain. All the intracranial injections were performed unilaterally.

For CTB experiments (Figure 1, 3, 4, 5), animals were sacrificed two days post-surgically. For monosynaptic rabies experiment (Figure 6), helper virus (AAV8-CAG-FLEX-TCB and AAV8-CAG-FLEX-oG) and AAVretro-cre-eBFP were delivered simultaneously four weeks prior to the rabies injection, and mice were sacrificed after eight-days following rabies injection. For triple tracing experiment (Figure 7), mice were allowed to survive for two weeks after viral injections.

### Immunohistochemistry

We deeply anesthetized animals by intraperitoneal injecting pentobarbital sodium solution (50 mg/kg) and perfused transcardially with saline, followed by 4% paraformaldehyde (PFA)-0.1 M phosphate buffer (PB, pH 7.4). Brains were freshly removed and post-fixed in 4% PFA-0.1 M PB for 2 hours at room temperature. Fixed brains were placed in 10% sucrose overnight at 4 °C and were embedded in 12% gelatin-10% sucrose. After fixation in 10% formalin-30% sucrose overnight at 4 °C, serial coronal sections were sliced with microtone (SM2000R, Leica) at 50 μm and collected in 0.1 M PB.

For CTB immunohistochemistry, slices were incubated in goat anti-cholera toxin B subunit primary antibody (1:15000, List labs, 703) over night at 4 °C and biotinylated horse anti-goat secondary antibody (1:2000, Vector, BA-9500) for 2h at room temperature. Slices were incubated with ABC complex (volume ratio 1:1, Vector, AK-5200) for 1.5 hours at room temperature. Next, labelling was visualized with DAB staining (1:150), and slices were mounted for light microscopy. For GFP_sm_-myc signal visualization, slices were incubated in goat anti-myc primary antibody (1:10000, Abcam, ab9132) over night at 4 °C, followed with Cy™5 donkey anti-goat secondary antibody (1:400, Jackson, 705-175-147) at room temperature. All slices for fluorescent microscopy were counterstained with DAPI. All antibodies were diluted in 2% normal horse serum-0.4% triton-0.1M PBS solution.

### Microscopy and Analysis

All overview sections were imaged with either a brightfield microscope (Nanozoomer 2.0-RS, Hamamatsu) or a fluorescent microscope (Axio Imager 2, ZEISS), and higher-magnification images were captured with a confocal microscope (LSM 700, ZEISS) for further quantification. For plotting the brain contours and labelling, we either imported brain images (from fluorescent sections) or used an Olympus microscope equipped with a Lucivid miniature monitor and manually plotted with Neurolucida software (Microbrightfield, VT, USA). We used serial sections with 100 μm intervals to plot as well as quantify labelling in the CN, the IO and the MDJ areas. Serial sections with 200 μm intervals were used for quantification and plotting of cortical neurons.

For the retrieval analysis of cortical injections from the Allen Mouse Brain, injection radius was firstly calculated by the formula: 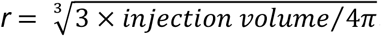, then we used the area calculated with *r* value to represent each injection and plotted on a standard flattened cortical map based on the given stereotaxic coordinates. For optimal visualization, we multiplied area values by 10 and transformed them into sized markers: circles for wildtype mice and squares for transgenic mice. To map the MDJ projection density on the cortical map (Figure 2C), we subjectively assessed cortical axon density on four coronal sections from the anteroposterior MDJ regions by scoring 0-3 for each section, in total 0-12 for each mouse. All injections were color coded based on the sum density scores. To plot MDJ-projection organisation on the cortical map (Figure 2B), we first removed the grey markers (no cortical axon observed), then colored each case based on the relative anteroposterior location of the MDJ sections that exhibiting most abundant cortical projection (magenta for rostral, mix for intermediate and cyan for caudal MDJ projection).

Standard flattened maps of the CC, the CN and the IO were made by transforming the mouse brain atlas (Paxinos and Franklin, 2019), which was described in detail in our previous work (Aoki et al., 2019, Suzuki et al., 2012, Ruigrok and Voogd, 2000). To present CTB- and rabies-labelled cells on color-coded flatten maps of the CC and the CN, we first reconstructed the contours of these structures and plotted the labelled cells, then superimposed sections with 200 μm bins for the CC and 100 μm bins for the CN, respectively. Retrogradely-labelled cells were counted in each bin, resulting in numeric matrices for the MDJ-projection cells in the CC and CN. For the IO colormap, we first marked the border of MDJ axon on each olivary section; next, we subjectively scored the axon density as 0-3 and illustrated on the standard flatten map of the IO. All colormaps were plotted using MATLAB.

To quantify the fraction of olivary-projecting MDJ cells (GFP) overlapping with CC and CN axons (Figure 7C), we applied thresholding (85th-95th percentiles) to binarize these images and then overlayed GFP channel with the other two channels to calculation the colocalization and overlapping area fraction. To examine the CC/CN terminations on the olivary-projecting MDJ neurons (Figure 7D), we acquired z-stack confocal images (0.35 μm sectioning) for the MDJ regions. Synaptic contacts were defined as colocalizations from orthogonal views at more than three different z planes. To quantify the CC/CN bouton sizes (Figure 7E), we first binarized and applied threshold (85th-95th percentiles) to the maximum intensity projection images of these boutons; next, bouton areas (μm^2^) were calculated by summing the bright pixels and transformed based on the image magnification. All image quantification was performed by using ImageJ. Statistics was performed by using GraphPad Prism, and statistical significance was defined as *P* < 0.05.

## Funding

This work was supported by ERC advanced, ZonMw, NWO-ALW, Medical NeuroDelta, INTENSE (LSH-NWO), Vriendenfonds Albinisme, and BIG (C.I.D.Z.); EUR fellowship, Erasmus MC fellowship, NWO VIDI, NWO-Klein and ERC-stg grants (Z.G.).

## Acknowledgements

We thank E.H. Sabel-Goedknegt for expert technical assistance; M. Rutteman for managing the animal breeding.

## Competing interests

We declare no financial or non-financial competing interests.

## Supplementary information

**Supplementary Table 1.** Stereotaxic coordinates of nuclei.

| Nuclei | AP (mm) | ML (mm) | Depth (mm) |
| --- | --- | --- | --- |
| Mesodiencephalic junction | -2.7 ~ -2.9 | 0.3 | 3.8 ~ 4.0 |
| Motor cortex | 1.2 | 1.0 | 0.6 |
| Periaqueductal grey | 2.8 | 0.0 | 3.8 |
| Inferior olive complex* | -2.8 | 0.3 ~ 0.5 | 5.4 |
| Cerebellar nuclei (interposed-dentate)* | 2.2 | 2.3 | 2.4 |
| Medullary gigantocellular reticular nucleus* | -2.8 | 0.5 | 4.9 |

\*The origin of coordinates is defined as the anterior tip of the interparietal bone; otherwise, coordinates were measured from the Bregma.
Viral/tracer injection volume was 20-50 nL. AP: anterior posterior, ML: medial lateral.

**Figure 1- figure supplement 1.**
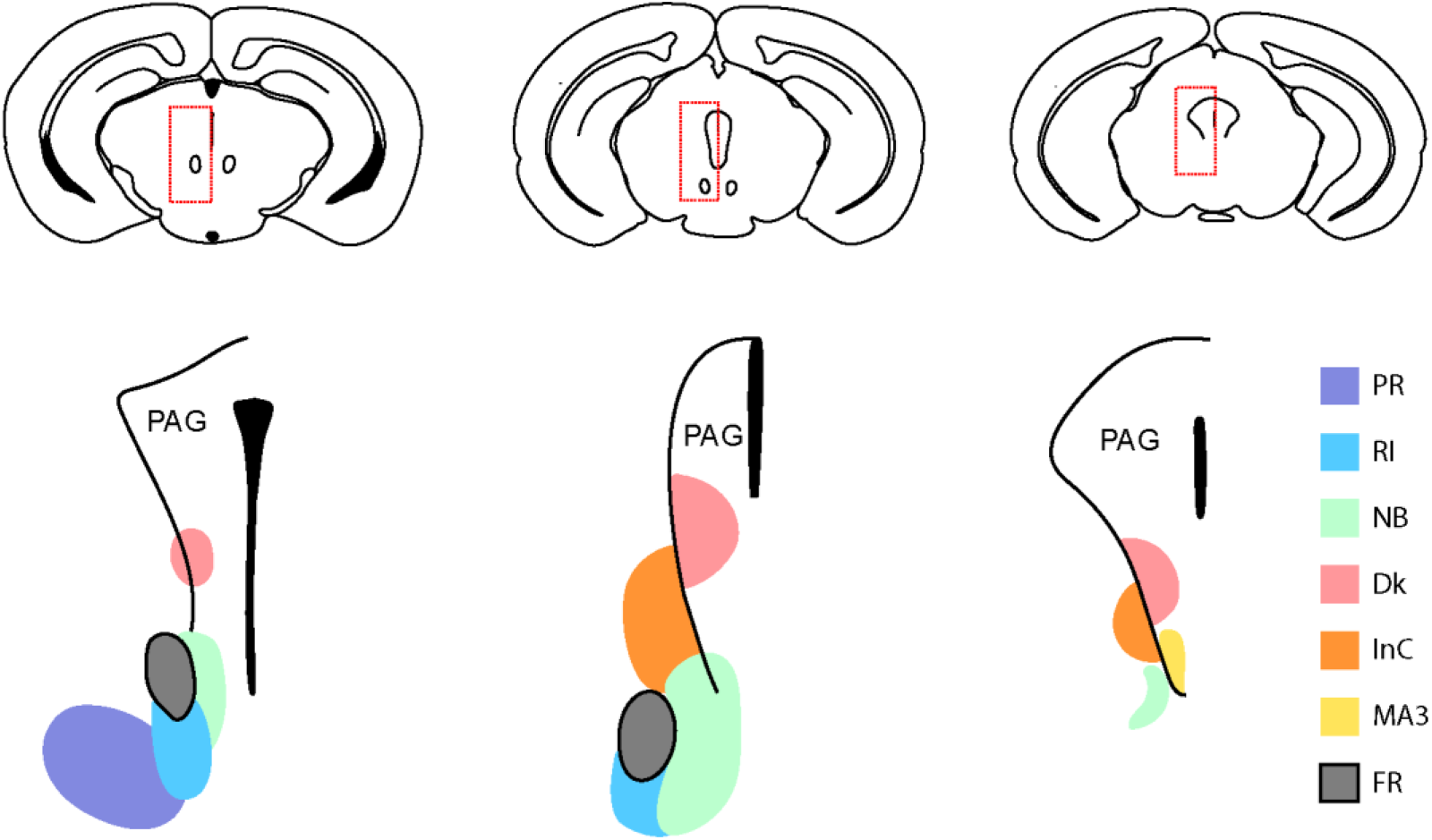
Architecture of the MDJ nuclei projecting to the IO. Areas on the left and right side represent the more rostral and caudal subregions of the MDJ, respectively. Abbreviations: PR: prerubral field, RI: rostral interstitial nucleus of the medial longitudinal fasciculus, NB: medial accessory nucleus of Bechterew, Dk: nucleus of Darkschewitsch, InC: interstitial nucleus of Cajal, MA3: medial accessory oculomotor nucleus, FR: fasciculus retroflexus.

**Figure 1 - figure supplement 2.**
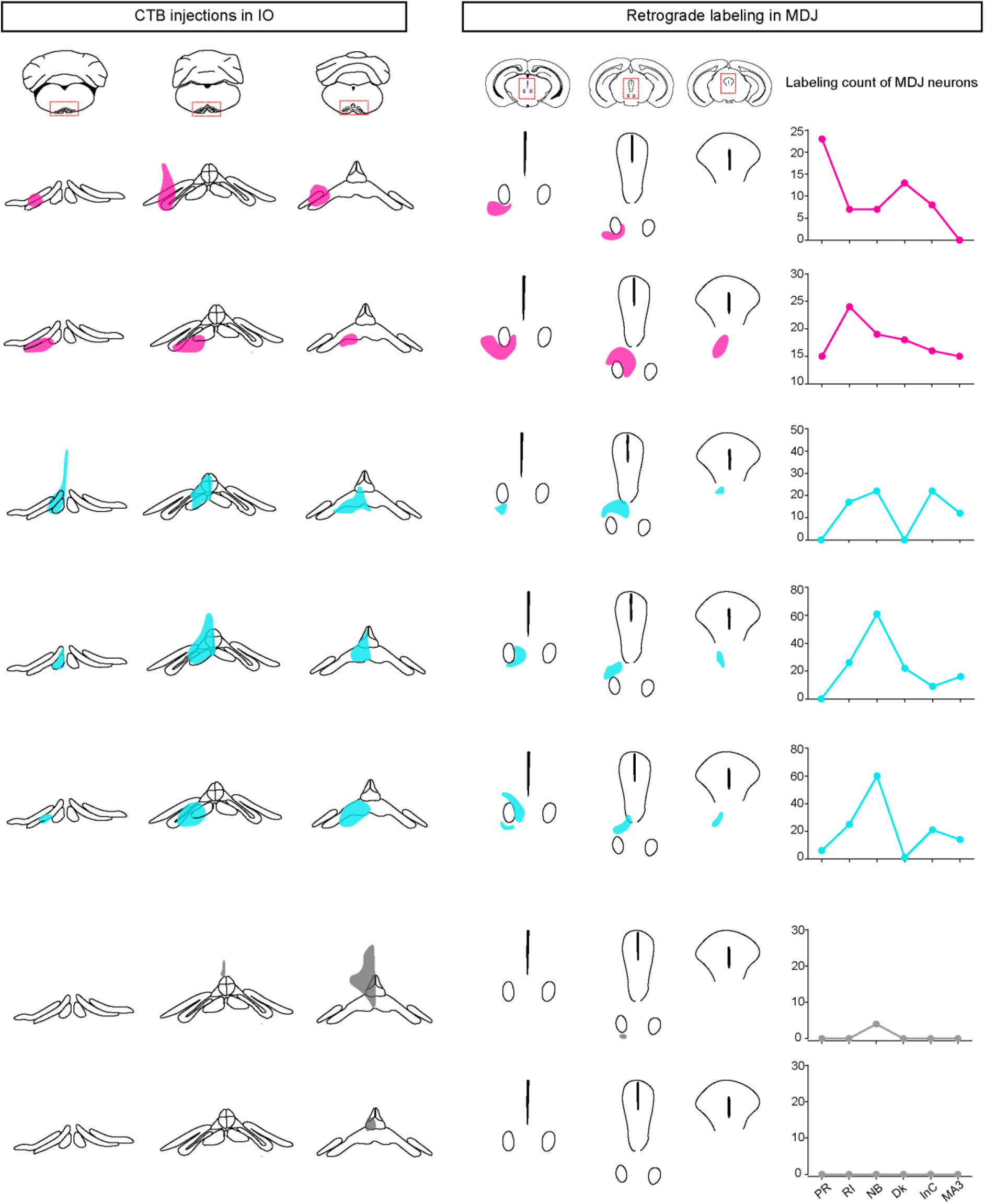
Individual CTB injections in the IO and retrograde labelling in the MDJ. Seven mice were injected with CTB in different subnuclei of the IO providing different sets of retrogradely labelled neurons in the MDJ at the various rostrocaudal levels. The injections are ordered according to their densities in the MDJ with the most prominent labelling in the rostral and caudal subregions at the top and bottom, respectively. Likewise, the MDJ subregions in the right column are ordered according to the rostrocaudal level of their core part. See abbreviations in **Figure 1-figure supplement 1**.

**Figure 1 - figure supplement 3.**
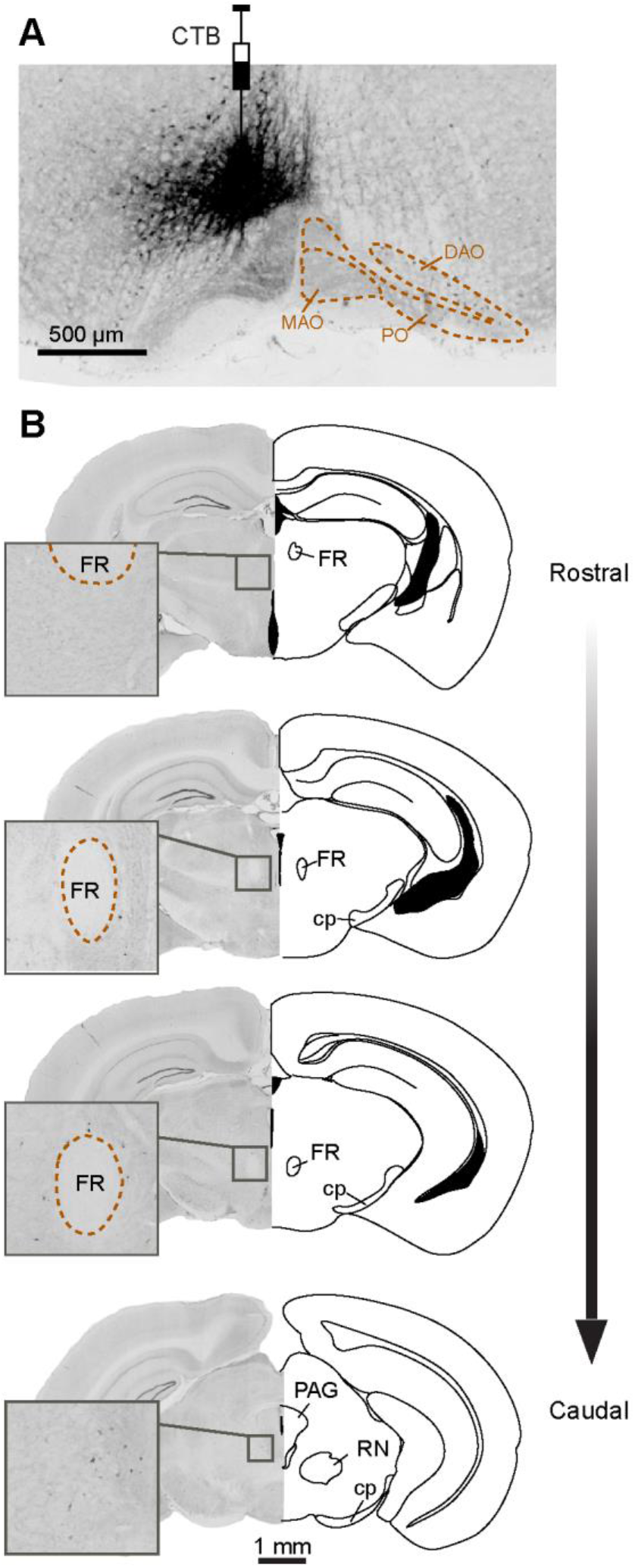
Retrogradely labelled cells in the MDJ following CTB injection in the medullary reticular formation. (A) Coronal section of the injection site from an example mouse. (B) Serial sections of anteroposterior MDJ regions, showing sparse retrograde labelling in the ipsilateral MDJ (inserts).

**Figure 3 - figure supplement 1.**
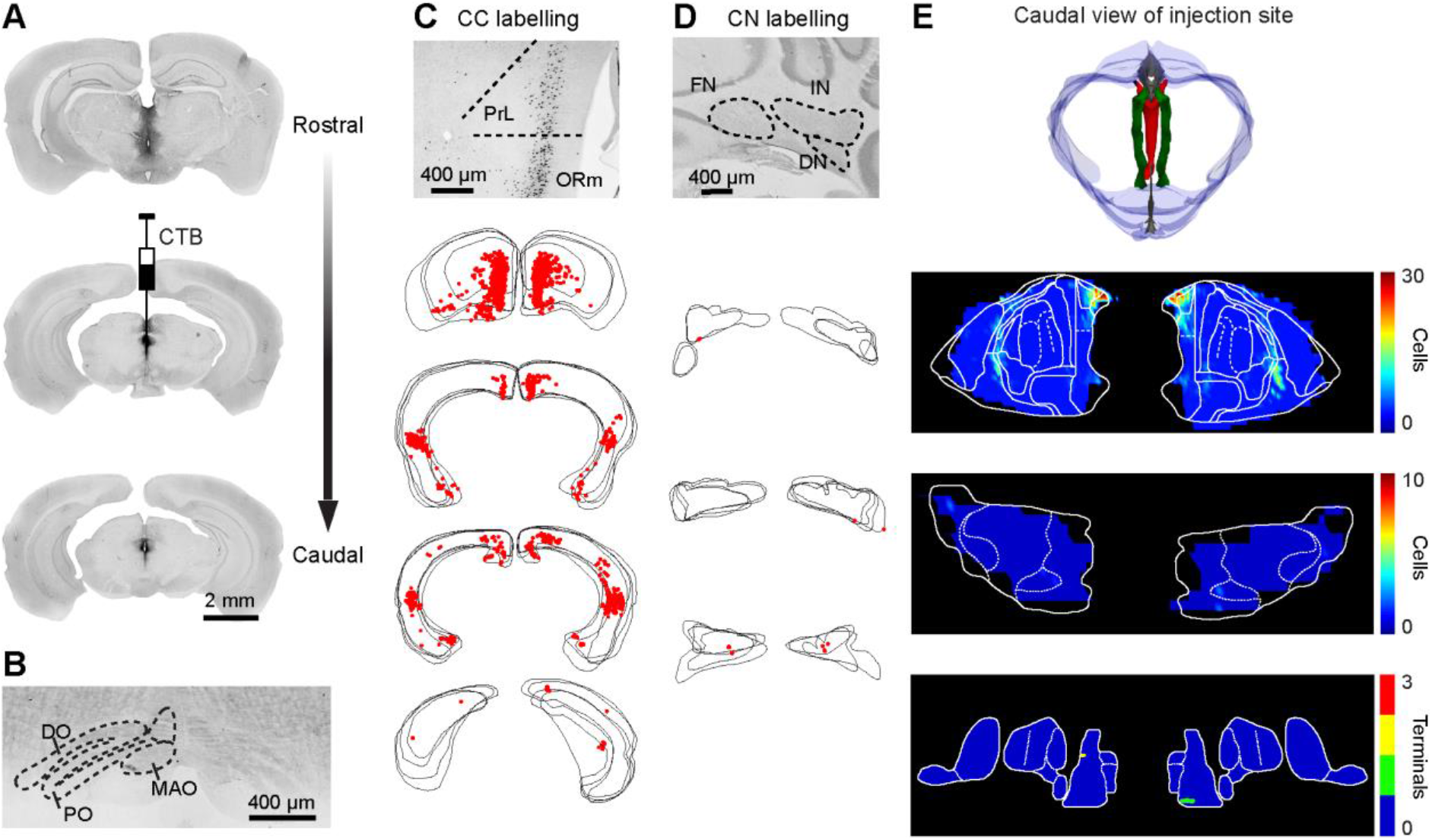
Input and output maps of the PAG. (A and B) Coronal sections showing CTB injection in the PAG (A) labelled sparse axons in the IO (B). (C and D) Retrogradely labelled cells in the CC and CN. Upper: representative sections showing dense CTB-labelled cells in the ipsilateral prefrontal cortical regions whereas few labelling in the contralateral CN. Lower: series plotting at longitudinal cortical areas and CN. (E) Summary of injection site and input, output maps of PAG. In the three-dimensional reconstruction of injection site, red: CTB injection site, blue: midbrain contour, grey: third ventricle, green: FR. See abbreviations in **Figure 2**.

## Notes

### Competing Interest Statement

The authors have declared no competing interest.

## References

Abdelgabar, A. R., Suttrup, J., Broersen, R., Bhandari, R., Picard, S., Keysers, C., De Zeeuw, C. I. & Gazzola, V. 2019. Action perception recruits the cerebellum and is impaired in patients with spinocerebellar ataxia. Brain, 142, 3791–3805.

Ackerley, R., Pardoe, J. & Apps, R. 2006. A novel site of synaptic relay for climbing fibre pathways relaying signals from the motor cortex to the cerebellar cortical C1 zone. J Physiol, 576, 503–18.

Akkal, D., Dum, R. P. & Strick, P. L. 2007. Supplementary motor area and presupplementary motor area: targets of basal ganglia and cerebellar output. J Neurosci, 27, 10659–73.

Amino, Y., Kyuhou, S., Matsuzaki, R. & Gemba, H. 2001. Cerebello-thalamo-cortical projections to the posterior parietal cortex in the macaque monkey. Neurosci Lett, 309, 29–32.

An, X., Bandler, R., OngÜr, D. & Price, J. L. 1998. Prefrontal cortical projections to longitudinal columns in the midbrain periaqueductal gray in macaque monkeys. J Comp Neurol, 401, 455–79.

Aoki, S., Coulon, P. & Ruigrok, T. J. H. 2019. Multizonal Cerebellar Influence Over Sensorimotor Areas of the Rat Cerebral Cortex. Cereb Cortex, 29, 598–614.

Apps, R. & Hawkes, R. 2009. Cerebellar cortical organization: a one-map hypothesis. Nat Rev Neurosci, 10, 670–81.

Barmack, N. H. 2006. Inferior olive and oculomotor system. Prog Brain Res, 151, 269–91.

Bentivoglio, M. & Kuypers, H. G. 1982. Divergent axon collaterals from rat cerebellar nuclei to diencephalon, mesencephalon, medulla oblongata and cervical cord. A fluorescent double retrograde labeling study. Exp Brain Res, 46, 339–56.

Berkley, K. J., Budell, R. J., Blomqvist, A. & Bull, M. 1986. Output systems of the dorsal column nuclei in the cat. Brain Res, 396, 199–225.

Berkley, K. J. & Worden, I. G. 1978. Projections to the inferior olive of the cat. I. Comparisons of input from the dorsal column nuclei, the lateral cervical nucleus, the spino-olivary pathways, the cerebral cortex and the cerebellum. J Comp Neurol, 180, 237–51.

Berretta, S., Bosco, G., Giaquinta, G., Smecca, G. & Perciavalle, V. 1993. Cerebellar influences on accessory oculomotor nuclei of the rat: a neuroanatomical, immunohistochemical, and electrophysiological study. J Comp Neurol, 338, 50–66.

Biswas, M. S., Luo, Y., Sarpong, G. A. & Sugihara, I. 2019. Divergent projections of single pontocerebellar axons to multiple cerebellar lobules in the mouse. J Comp Neurol, 527, 1966–1985.

Braak, H., RÜb, U. & Del Tredici, K. 2003. Involvement of precerebellar nuclei in multiple system atrophy. Neuropathol Appl Neurobiol, 29, 60–76.

Brissenden, J. A. & Somers, D. C. 2019. Cortico-cerebellar networks for visual attention and working memory. Curr Opin Psychol, 29, 239–247.

Brodal, P. 1978. Principles of organization of the monkey corticopontine projection. Brain Res, 148, 214–8.

Brown, J. T., Chan-Palay, V. & Palay, S. L. 1977. A study of afferent input to the inferior olivary complex in the rat by retrograde axonal transport of horseradish peroxidase. J Comp Neurol, 176, 1–22.

Brown, S. T. & Raman, I. M. 2018. Sensorimotor Integration and Amplification of Reflexive Whisking by Well-Timed Spiking in the Cerebellar Corticonuclear Circuit. Neuron, 99, 564–575 e2.

Bull, M. S., Mitchell, S. K. & Berkley, K. J. 1990. Convergent inputs to the inferior olive from the dorsal column nuclei and pretectum in the cat. Brain Res, 525, 1–10.

Burman, K., Darian-Smith, C. & Darian-Smith, I. 2000. Macaque red nucleus: origins of spinal and olivary projections and terminations of cortical inputs. J Comp Neurol, 423, 179–96.

Callaway, E. M. & Luo, L. 2015. Monosynaptic Circuit Tracing with Glycoprotein-Deleted Rabies Viruses. The Journal of neuroscience: the official journal of the Society for Neuroscience, 35, 8979–8985.

Carlton, S. M., Leichnetz, G. R. & Mayer, J. D. 1982. Projections from the nucleus parafascicularis prerubralis to medullary raphe nuclei and inferior olive in the rat: a horseradish peroxidase and autoradiography study. Neurosci Lett, 30, 191–7.

Chabrol, F. P., Blot, A. & Mrsic-Flogel, T. D. 2019. Cerebellar Contribution to Preparatory Activity in Motor Neocortex. Neuron, 103, 506–519 e4.

Coffman, K. A., Dum, R. P. & Strick, P. L. 2011. Cerebellar vermis is a target of projections from the motor areas in the cerebral cortex. Proc Natl Acad Sci U S A, 108, 16068–73.

CondÉ, F. & CondÉ, H. 1982. The rubro-olivary tract in the cat, as demonstrated with the method of retrograde transport of horseradish peroxidase. Neuroscience, 7, 715–24.

Cornwall, J. & Phillipson, O. T. 1988. Afferent projections to the parafascicular thalamic nucleus of the rat, as shown by the retrograde transport of wheat germ agglutinin. Brain Res Bull, 20, 139–50.

Crill, W. E. 1970. Unitary multiple-spiked responses in cat inferior olive nucleus. J Neurophysiol, 33, 199–209.

Crippa, A., Del Vecchio, G., Busti Ceccarelli, S., Nobile, M., Arrigoni, F. & Brambilla, P. 2016. Cortico-Cerebellar Connectivity in Autism Spectrum Disorder: What Do We Know So Far? Front Psychiatry, 7, 20.

D’mello, A. M. & Stoodley, C. J. 2015. Cerebro-cerebellar circuits in autism spectrum disorder. Front Neurosci, 9, 408.

DE ZEEUW, C. I. 2021. Bidirectional learning in upbound and downbound microzones of the cerebellum. Nat Rev Neurosci, 22, 92–110.

De Zeeuw, C. I., Holstege, J. C., Ruigrok, T. J. & Voogd, J. 1989. Ultrastructural study of the GABAergic, cerebellar, and mesodiencephalic innervation of the cat medial accessory olive: anterograde tracing combined with immunocytochemistry. J Comp Neurol, 284, 12–35.

De Zeeuw, C. I. & Ruigrok, T. J. 1994. Olivary projecting neurons in the nucleus of Darkschewitsch in the cat receive excitatory monosynaptic input from the cerebellar nuclei. Brain Res, 653, 345–50.

De Zeeuw, C. I., Simpson, J. I., Hoogenraad, C. C., Galjart, N., Koekkoek, S. K. & Ruigrok, T. J. 1998. Microcircuitry and function of the inferior olive. Trends Neurosci, 21, 391–400.

Depping, M. S., Schmitgen, M. M., Kubera, K. M. & Wolf, R. C. 2018. Cerebellar Contributions to Major Depression. Front Psychiatry, 9, 634.

Fujita, H., Kodama, T. & Du Lac, S. 2020. Modular output circuits of the fastigial nucleus for diverse motor and nonmotor functions of the cerebellar vermis. Elife, 9.

Fukushima, K., Terashima, T., Kudo, J., Inoue, Y. & Kato, M. 1986. Projections of the group y of the vestibular nuclei and the dentate and fastigial nuclei of the cerebellum to the interstitial nucleus of Cajal. Neurosci Res, 3, 285–99.

Gao, Z., Davis, C., Thomas, A. M., Economo, M. N., Abrego, A. M., Svoboda, K., De Zeeuw, C. I. & Li, N. 2018. A cortico-cerebellar loop for motor planning. Nature, 563, 113–116.

Glickstein, M., May, J. G. 3rd & Mercier, B. E. 1985. Corticopontine projection in the macaque: the distribution of labelled cortical cells after large injections of horseradish peroxidase in the pontine nuclei. J Comp Neurol, 235, 343–59.

Gonzalo-Ruiz, A., Leichnetz, G. R. & Hardy, S. G. 1990. Projections of the medial cerebellar nucleus to oculomotor-related midbrain areas in the rat: an anterograde and retrograde HRP study. J Comp Neurol, 296, 427–36.

Gonzalo-Ruiz, A., Leichnetz, G. R. & Smith, D. J. 1988. Origin of cerebellar projections to the region of the oculomotor complex, medial pontine reticular formation, and superior colliculus in New World monkeys: a retrograde horseradish peroxidase study. J Comp Neurol, 268, 508–26.

Guo, J.-Z., Sauerbrei, B., Cohen, J. D., Mischiati, M., Graves, A., Pisanello, F., Branson, K. & Hantman, A. W. 2020. Dynamics of the Cortico-Cerebellar Loop Fine-Tune Dexterous Movement. bioRxiv, 637447.

Halmagyi, G. M., Aw, S. T., Dehaene, I., Curthoys, I. S. & Todd, M. J. 1994. Jerk-waveform see-saw nystagmus due to unilateral meso-diencephalic lesion. Brain, 117 (Pt 4), 789–803.

Hamada, M., Strigaro, G., Murase, N., Sadnicka, A., Galea, J. M., Edwards, M. J. & Rothwell, J. C. 2012. Cerebellar modulation of human associative plasticity. J Physiol, 590, 2365–74.

Heffley, W., Song, E. Y., Xu, Z., Taylor, B. N., Hughes, M. A., Mckinney, A., Joshua, M. & Hull, C. 2018. Coordinated cerebellar climbing fiber activity signals learned sensorimotor predictions. Nat Neurosci, 21, 1431–1441.

Henschke, J. U. & Pakan, J. M. 2020. Disynaptic cerebrocerebellar pathways originating from multiple functionally distinct cortical areas. Elife, 9.

Hoover, W. B. & Vertes, R. P. 2007. Anatomical analysis of afferent projections to the medial prefrontal cortex in the rat. Brain Struct Funct, 212, 149–79.

IgelstrÖm, K. M., Webb, T. W. & Graziano, M. S. A. 2017. Functional Connectivity Between the Temporoparietal Cortex and Cerebellum in Autism Spectrum Disorder. Cereb Cortex, 27, 2617–2627.

Kalil, K. 1981. Projections of the cerebellar and dorsal column nuclei upon the thalamus of the rhesus monkey. J Comp Neurol, 195, 25–50.

Kato, N., Kawaguchi, S. & Miyata, H. 1988. Cerebro-cerebellar projections from the lateral suprasylvian visual area in the cat. J Physiol, 395, 473–85.

Kim, E. J., Jacobs, M. W., Ito-Cole, T. & Callaway, E. M. 2016. Improved Monosynaptic Neural Circuit Tracing Using Engineered Rabies Virus Glycoproteins. Cell reports, 15, 692–699.

Kostadinov, D., Beau, M., Blanco-Pozo, M. & HÄusser, M. 2019. Predictive and reactive reward signals conveyed by climbing fiber inputs to cerebellar Purkinje cells. Nat Neurosci, 22, 950–962.

Kubo, R., Aiba, A. & Hashimoto, K. 2018. The anatomical pathway from the mesodiencephalic junction to the inferior olive relays perioral sensory signals to the cerebellum in the mouse. J Physiol, 596, 3775–3791.

Larson, B., Miller, S. & Oscarsson, O. 1969. Termination and functional organization of the dorsolateral spino-olivocerebellar path. J Physiol, 203, 611–40.

Leergaard, T. B. 2003. Clustered and laminar topographic patterns in rat cerebro-pontine pathways. Anat Embryol (Berl), 206, 149–62.

Leergaard, T. B., Lyngstad, K. A., Thompson, J. H., Taeymans, S., Vos, B. P., De Schutter, E., Bower, J. M. & Bjaalie, J. G. 2000. Rat somatosensory cerebropontocerebellar pathways: spatial relationships of the somatotopic map of the primary somatosensory cortex are preserved in a three-dimensional clustered pontine map. J Comp Neurol, 422, 246–66.

Legg, C. R., Mercier, B. & Glickstein, M. 1989. Corticopontine projection in the rat: the distribution of labelled cortical cells after large injections of horseradish peroxidase in the pontine nuclei. J Comp Neurol, 286, 427–41.

Leichnetz, G. R., Spencer, R. F. & Smith, D. J. 1984. Cortical projections to nuclei adjacent to the oculomotor complex in the medial dien-mesencephalic tegmentum in the monkey. J Comp Neurol, 228, 359–87.

Linauts, M. & Martin, G. F. 1978. An autoradiographic study of midbrain-diencephalic projections to the inferior olivary nucleus in the opossum (Didelphis virginiana). J Comp Neurol, 179, 325–53.

Lindeman, S., Hong, S., Kros, L., Mejias, J. F., Romano, V., Oostenveld, R., Negrello, M., Bosman, L. W. J. & De Zeeuw, C. I. 2021. Cerebellar Purkinje cells can differentially modulate coherence between sensory and motor cortex depending on region and behavior. Proc Natl Acad Sci U S A, 118.

Mabuchi, M. & Kusama, T. 1970. Mesodiencephalic projections to the inferior olive and the vestibular and perihypoglossal nuclei. Brain research, 17, 133–136.

Mandelbaum, G., Taranda, J., Haynes, T. M., Hochbaum, D. R., Huang, K. W., Hyun, M., Umadevi Venkataraju, K., Straub, C., Wang, W., Robertson, K., Osten, P. & Sabatini, B. L. 2019. Distinct Cortical-Thalamic-Striatal Circuits through the Parafascicular Nucleus. Neuron, 102, 636–652 e7.

Massion, J. 1967. The mammalian red nucleus. Physiol Rev, 47, 383–436.

Mccrea, R. A. & Baker, R. 1985. Anatomical connections of the nucleus prepositus of the cat. J Comp Neurol, 237, 377–407.

Mitrofanis, J. & Defonseka, R. 2001. Organisation of connections between the zona incerta and the interposed nucleus. Anat Embryol (Berl), 204, 153–9.

Murphy, M. J. M. & Deutch, A. Y. 2018. Organization of afferents to the orbitofrontal cortex in the rat. J Comp Neurol, 526, 1498–1526.

Nakamura, Y., Kitao, Y. & Okoyama, S. 1983a. Cortico-Darkschewitsch-olivary projection in the cat: an electron microscope study with the aid of horseradish peroxidase tracing technique. Brain Res, 274, 140–3.

Nakamura, Y., Kitao, Y. & Okoyama, S. 1983b. Projections from the pericruciate cortex to the nucleus of Darkschewitsch and other structures at the mesodiencephalic junction in the cat. Brain Res Bull, 10, 517–21.

Negrello, M., Warnaar, P., Romano, V., Owens, C. B., Lindeman, S., Iavarone, E., Spanke, J. K., Bosman, L. W. J. & De Zeeuw, C. I. 2019. Quasiperiodic rhythms of the inferior olive. PLoS Comput Biol, 15, e1006475.

Oh, S. W., Harris, J. A., Ng, L., Winslow, B., Cain, N., Mihalas, S., Wang, Q., Lau, C., Kuan, L., Henry, A. M., Mortrud, M. T., Ouellette, B., Nguyen, T. N., Sorensen, S. A., Slaughterbeck, C. R., Wakeman, W., Li, Y., Feng, D., Ho, A., Nicholas, E., Hirokawa, K. E., Bohn, P., Joines, K. M., Peng, H., Hawrylycz, M. J., Phillips, J. W., Hohmann, J. G., Wohnoutka, P., Gerfen, C. R., Koch, C., Bernard, A., Dang, C., Jones, A. R. & Zeng, H. 2014. A mesoscale connectome of the mouse brain. Nature, 508, 207–14.

Ohmae, S., Kunimatsu, J. & Tanaka, M. 2017. Cerebellar Roles in Self-Timing for Sub- and Supra-Second Intervals. J Neurosci, 37, 3511–3522.

Ohmae, S. & Medina, J. F. 2015. Climbing fibers encode a temporal-difference prediction error during cerebellar learning in mice. Nature Neuroscience, 18, 1798–1803.

Olivito, G., Clausi, S., Laghi, F., Tedesco, A. M., Baiocco, R., Mastropasqua, C., Molinari, M., Cercignani, M., Bozzali, M. & Leggio, M. 2017. Resting-State Functional Connectivity Changes Between Dentate Nucleus and Cortical Social Brain Regions in Autism Spectrum Disorders. Cerebellum, 16, 283–292.

Onodera, S. 1984. Olivary projections from the mesodiencephalic structures in the cat studied by means of axonal transport of horseradish peroxidase and tritiated amino acids. J Comp Neurol, 227, 37–49.

Onodera, S. & Hicks, T. P. 1996. A projection linking motor cortex with the LM-suprageniculate nuclear complex through the periaqueductal gray area which surrounds the nucleus of Darkschewitsch in the cat. Prog Brain Res, 112, 85–98.

Onodera, S. & Hicks, T. P. 2009. A comparative neuroanatomical study of the red nucleus of the cat, macaque and human. PLoS One, 4, e6623.

Onuki, Y., Van Someren, E. J., De Zeeuw, C. I. & Van Der Werf, Y. D. 2015. Hippocampal-cerebellar interaction during spatio-temporal prediction. Cereb Cortex, 25, 313–21.

Pardoe, J., Edgley, S. A., Drew, T. & Apps, R. 2004. Changes in excitability of ascending and descending inputs to cerebellar climbing fibers during locomotion. J Neurosci, 24, 2656–66.

Paxinos, G. & Franklin, K. B. 2019. Paxinos and Franklin’s the mouse brain in stereotaxic coordinates, Academic press.

Peschanski, M. & Mantyh, P. W. 1983. Efferent connections of the subfascicular area of the mesodiencephalic junction and its possible involvement in stimulation-produced analgesia. Brain Res, 263, 181–90.

Popa, T., Velayudhan, B., Hubsch, C., Pradeep, S., Roze, E., Vidailhet, M., Meunier, S. & Kishore, A. 2013. Cerebellar processing of sensory inputs primes motor cortex plasticity. Cereb Cortex, 23, 305–14.

Proville, R. D., Spolidoro, M., Guyon, N., DuguÉ, G. P., Selimi, F., Isope, P., Popa, D. & LÉna, C. 2014. Cerebellum involvement in cortical sensorimotor circuits for the control of voluntary movements. Nat Neurosci, 17, 1233–9.

Quartarone, A., Cacciola, A., Milardi, D., Ghilardi, M. F., Calamuneri, A., Chillemi, G., Anastasi, G. & Rothwell, J. 2020. New insights into cortico-basal-cerebellar connectome: clinical and physiological considerations. Brain, 143, 396–406.

Ramnani, N. 2006. The primate cortico-cerebellar system: anatomy and function. Nature Reviews Neuroscience, 7, 511–522.

Rispal-Padel, L. & Grangetto, A. 1977. The cerebello-thalamo-cortical pathway. Topographical investigation at the unitary level in the cat. Exp Brain Res, 28, 101–23.

Romano, V., De Propris, L., Bosman, L. W., Warnaar, P., Ten Brinke, M. M., Lindeman, S., Ju, C., Velauthapillai, A., Spanke, J. K., Middendorp Guerra, E., Hoogland, T. M., Negrello, M., D’angelo, E. & De Zeeuw, C. I. 2018. Potentiation of cerebellar Purkinje cells facilitates whisker reflex adaptation through increased simple spike activity. Elife, 7.

Royce, G. J., Bromley, S. & Gracco, C. 1991. Subcortical projections to the centromedian and parafascicular thalamic nuclei in the cat. J Comp Neurol, 306, 129–55.

Ruigrok, T. J. 2004. Precerebellar nuclei and red nucleus. The rat nervous system, 167–204.

Ruigrok, T. J. 2011. Ins and outs of cerebellar modules. Cerebellum, 10, 464–74.

Ruigrok, T. J. & Teune, T. M. 2014. Collateralization of cerebellar output to functionally distinct brainstem areas. A retrograde, non-fluorescent tracing study in the rat. Front Syst Neurosci, 8, 23.

Ruigrok, T. J. & Voogd, J. 1995. Cerebellar influence on olivary excitability in the cat. Eur J Neurosci, 7, 679–93.

Ruigrok, T. J. & Voogd, J. 2000. Organization of projections from the inferior olive to the cerebellar nuclei in the rat. J Comp Neurol, 426, 209–28.

Rutherford, J. G., Anderson, W. A. & Gwyn, D. G. 1984. A reevaluation of midbrain and diencephalic projections to the inferior olive in rat with particular reference to the rubro-olivary pathway. J Comp Neurol, 229, 285–300.

Rutherford, J. G., Zuk-Harper, A. & Gwyn, D. G. 1989. A comparison of the distribution of the cerebellar and cortical connections of the nucleus of Darkschewitsch (ND) in the cat: a study using anterograde and retrograde HRP tracing techniques. Anat Embryol (Berl), 180, 485–96.

Saint-Cyr, J. A. 1983. The projection from the motor cortex to the inferior olive in the cat. An experimental study using axonal transport techniques. Neuroscience, 10, 667–84.

Saint-Cyr, J. A. 1987. Anatomical organization of cortico-mesencephalo-olivary pathways in the cat as demonstrated by axonal transport techniques. J Comp Neurol, 257, 39–59.

Saint-Cyr, J. A. & Courville, J. 1981. Sources of descending afferents to the inferior olive from the upper brain stem in the cat as revealed by the retrograde transport of horseradish peroxidase. J Comp Neurol, 198, 567–81.

Sasaki, K., Oka, H., Matsuda, Y., Shimono, T. & Mizuno, N. 1975. Electrophysiological studies of the projections from the parietal association area to the cerebellar cortex. Exp Brain Res, 23, 91–102.

Shinoda, Y., Kakei, S., Futami, T. & Wannier, T. 1993. Thalamocortical organization in the cerebello-thalamo-cortical system. Cereb Cortex, 3, 421–9.

Shiraishi, Y. & Nakao, S. 1994. Projections of vertical eye movement-related and head rotation-related neurons in the medial mesodiencephalic junction to pontine reticular formation in cat. Neurosci Lett, 171, 85–8.

Stanton, G. B. 1980. Topographical organization of ascending cerebellar projections from the dentate and interposed nuclei in Macaca mulatta: an anterograde degeneration study. J Comp Neurol, 190, 699–731.

Strominger, N. L., Truscott, T. C., Miller, R. A. & Royce, G. J. 1979. An autoradiographic study of the rubroolivary tract in the rhesus monkey. J Comp Neurol, 183, 33–45.

Suzuki, L., Coulon, P., Sabel-Goedknegt, E. H. & Ruigrok, T. J. 2012. Organization of cerebral projections to identified cerebellar zones in the posterior cerebellum of the rat. J Neurosci, 32, 10854–69.

Swenson, R. S. & Castro, A. J. 1983. The afferent connections of the inferior olivary complex in rats. An anterograde study using autoradiographic and axonal degeneration techniques. Neuroscience, 8, 259–275.

Ten Brinke, M. M., Boele, H.-J., Spanke, J. K., Potters, J.-W., Kornysheva, K., Wulff, P., Ijpelaar, A. C. H. G., Koekkoek, S. K. E. & De Zeeuw, C. I. 2015. Evolving Models of Pavlovian Conditioning: Cerebellar Cortical Dynamics in Awake Behaving Mice. Cell Reports, 13, 1977–1988.

Teune, T. M., Van Der Burg, J., Van Der Moer, J., Voogd, J. & Ruigrok, T. J. 2000. Topography of cerebellar nuclear projections to the brain stem in the rat. Prog Brain Res, 124, 141–72.

Voges, K., Wu, B., Post, L., Schonewille, M. & De Zeeuw, C. I. 2017. Mechanisms underlying vestibulo-cerebellar motor learning in mice depend on movement direction. J Physiol, 595, 5301–5326.

Voogd, J. & Ruigrok, T. J. 2004. The organization of the corticonuclear and olivocerebellar climbing fiber projections to the rat cerebellar vermis: the congruence of projection zones and the zebrin pattern. J Neurocytol, 33, 5–21.

Wagner, M. J., Kim, T. H., Kadmon, J., Nguyen, N. D., Ganguli, S., Schnitzer, M. J. & Luo, L. 2019. Shared Cortex-Cerebellum Dynamics in the Execution and Learning of a Motor Task. Cell, 177, 669–682 e24.

Walberg, F. & Nordby, T. 1981. A re-examination of the rubro-olivary tract in the cat, using horseradish peroxidase as a retrograde and an anterograde neuronal tracer. Neuroscience, 6, 2379–91.

Wall, N. R., Wickersham, I. R., Cetin, A., De La Parra, M. & Callaway, E. M. 2010. Monosynaptic circuit tracing in vivo through Cre-dependent targeting and complementation of modified rabies virus. Proceedings of the National Academy of Sciences of the United States of America, 107, 21848–21853.

Watson, T., Jones, M. & Apps, R. 2009. Electrophysiological mapping of novel prefrontal - cerebellar pathways. Frontiers in Integrative Neuroscience, 3.

Wiesendanger, R. & Wiesendanger, M. 1982. The corticopontine system in the rat. II. The projection pattern. J Comp Neurol, 208, 227–38.

Wiest, G., Baumgartner, C., Schnider, P., Trattnig, S., Deecke, L. & Mueller, C. 1996. Monocular elevation paresis and contralateral downgaze paresis from unilateral mesodiencephalic infarction. J Neurol Neurosurg Psychiatry, 60, 579–81.

